# FOXP-stabilization of the *Il2ra* super-enhancer structure augments Treg fitness

**DOI:** 10.64898/2026.04.14.718220

**Authors:** Dachuan Dong, Lauren E. Higdon, Jiayan Zhou, Jian-Xin Lin, Jyothi Padiapu, Yesl Kim, Warren J. Leonard, Jonathan S. Maltzman

## Abstract

Gene expression in regulatory T cells (Tregs) is context-dependent and maintains peripheral immune homeostasis. FOXP3 is lineage defining but not sufficient for Treg function or persistence. To define the cell-intrinsic roles of the FOXP3 paralogs FOXP1 and FOXP4, we generated and studied mice with Treg-specific deletion of *Foxp1* and/or *Foxp4*. FOXP1 and FOXP4 are required to maintain the peripheral Treg pool through enhancing *Il2ra* transcription, thereby promoting sustained high-level expression of IL-2Rα and thus of the high-affinity IL-2Rαβγ complex. Integrating RNA-seq and ATAC-seq with previously published ChIA-PET and publicly available data, we propose a model of *Il2ra* transcriptional regulation in which in which FOXP1 and FOXP4 anchor chromatin looping of the *Il2ra* locus in mature Tregs, augment super-enhancer activity, and drive sustained CD25 expression. Our results reveal a unique role of FOXP1, and to a lesser extent FOXP4, in controlling Treg homeostasis.

**One Sentence Summary:** FOXP1 and FOXP4 regulate chromatin architecture at the *Il2ra* locus, promoting sustained CD25 expression and maintaining the peripheral Treg pool.

## INTRODUCTION

CD4^+^FOXP3^+^ regulatory T cells (Tregs) are critical for maintaining immune homeostasis and self-tolerance. The X-chromosome encoded transcriptional factor Forkhead Box Protein P3 (FOXP3), also known as Scurfin, has broadly been recognized as the Treg lineage-defining factor in both humans and rodents(*1–3*). In murine cells expressing a *Foxp3* reporter null allele (Treg “wannabe”), the majority of loci with FOXP3-dependent changes are not directly bound by FOXP3(*4*). Chemogenetic degradation of FOXP3 in mature Treg has minimal impact on FOXP3 targeted genes under steady state conditions(*5*). These finding suggest that FOXP3 functions in a partially indirect manner by fine-tuning intermediates and by interacting with other transcriptional regulators. Notably, FOXP3 preferentially binds to already established active enhancers, and its transcriptional activity is fine-tuned in a partner-dependent manner(*6*). The molecular details of the factors that FOXP3 binds and how FOXP3 alone or with these additional factors regulates transcription have not been fully elucidated.

FOXP3 has three paralogs; FOXP1 and FOXP4 are also expressed in T lymphocytes, whereas FOXP2 is not. Both FOXP1 and FOXP4 have been identified as components of the FOXP3 supramolecular complex in murine Tregs(*7*) and the human Jurkat T cell line(*8*). FOXP1 is essential for maintaining quiescence of naïve T cells (Tnv), ensuring an appropriate germinal center response, and regulating cell death in naïve and effector T cell subsets(*9–11*). In Tregs, FOXP1 is necessary for facilitating FOXP3 binding to DNA(*12*) and optimum expression of FOXP3 in iTregs(*13*). In contrast, the role of FOXP4 in Tregs remains unclear. Our previous work has shown that Tregs can develop normally in the absence of FOXP4(*14*) but that the combined absence of FOXP1 and FOXP4 results in an activated/effector-like phenotype with compromised suppressive function, resulting in profound lymphoproliferation, inflammation, and early lethality(*15*).

Most of the above observations were made in mice with inflammatory microenvironments that precluded the ability to rigorously study and understand cell-intrinsic mechanisms. Here, we aimed to better understand the cell-intrinsic roles of FOXP1 and FOXP4, including their underlying transcriptional and epigenetic activities. Loss of FOXP1/FOXP4 reduces expression of the high-affinity interleukin-2 receptor (IL-2Rαβγ)(*16*) complex, which limits responsiveness to exogenous IL-2 in mature Tregs. Consequently, under competitive homeostatic conditions, FOXP1/FOXP4-deficient Tregs exhibit reduced fitness and are at a competitive disadvantage relative to FOXP1/FOXP4-sufficient Tregs. We further suggest an essential role of FOXP proteins in stabilizing intronic super-enhancer looping at the *Il2ra* locus.

## RESULTS

### Loss of FOXP1/FOXP4 reduces Treg cell fitness

Our previous results showed that mice with conditional deletion of FOXP1/FOXP4 in FOXP3^+^ Tregs developed a fatal autoinflammatory phenotype(*15*). To exclude the impact of inflammation-associated cell-extrinsic factors, here we used *Foxp3*^YFP-Cre/X^ heterozygous deleters. We crossed hemizygous males (*Foxp3*^YFP-Cre/Y^) with wild-type females (*Foxp3*^X/X^) to generate *Foxp3*^YFP-Cre/X^ heterozygous female offspring (fig. S1A), which exhibit chimerism of Tregs due to random X-inactivation. Individual Tregs either express the YFP reporter and Cre recombinase to excise any *loxP* flanked sites or they lack Cre and maintain a YFP^-^ phenotype. At postnatal day 21, we weaned and co-housed *Foxp3*^YFP-Cre/X^*Foxp1/4*^fl/fl^ mice with age-matched *Foxp3*^YFP-Cre/X^*Foxp1/4*^+/+^ controls. At 8-weeks of age, the splenomegaly and lymphoproliferation observed in *Foxp3*^YFP-Cre/YFP-Cre^*Foxp1/4*^fl/fl^ homozygous females were absent, suggesting that Cre-negative, FOXP1/FOXP4-sufficient Tregs were capable of maintaining peripheral immune homeostasis despite the presence of FOXP1/FOXP4 deficient Tregs (fig. S1B).

We first evaluated the impact that loss of FOXP1 and FOXP4 has on competitive fitness within the FOXP3^+^ compartment by comparing the relative frequency of YFP^+^ versus YFP^-^ cells. Consistent with the competitive disadvantage of FOXP1-deficient Tregs previously observed in the secondary lymphoid organs (SLOs) by others(*12*), we noted that the loss of FOXP1 alone (*Foxp3*^YFP-Cre/X^*Foxp1*^fl/fl^*Foxp4*^+/+^) or combined with loss of FOXP4 (*Foxp3*^YFP-Cre/X^*Foxp1/4*^fl/fl^), resulted in a decrease in Cre-expressing YFP^+^ Tregs and a concomitant increase in YFP^-^ Tregs that lack Cre-mediated deletion (Fig. 1A). In contrast, the absence of FOXP4 alone (*Foxp3*^YFP-Cre/X^*Foxp1*^+/+^*Foxp4*^fl/fl^) had a negligible impact on Treg competitiveness in SLOs (Fig. 1A). To avoid potential compensatory effects from FOXP4, we primarily focused on *Foxp3^YFP-^*^Cre/X^*Foxp1/4*^fl/fl^ mice (*Foxp3* persistent). To determine if the competitive disadvantage caused by loss of FOXP1 and FOXP4 had a temporal component, we conducted a longitudinal study. The lack of FOXP1 and FOXP4 had no consistent effect on the total number of peripheral Tregs nor the relative proportion of Tregs within the peripheral CD4 compartment except for a transient difference at 4 weeks of age in the spleen (Fig. 1B). By postnatal week 4, both the cellularity and proportion of YFP^+^ Treg cells had declined in *Foxp3*^YFP*-*Cre/X^*Foxp1/4*^fl/fl^ mice relative to *Foxp3*^YFP*-*Cre/X^*Foxp1/4*^+/+^ mice (Fig. 1C and fig. S1C).

**Fig. 1.**
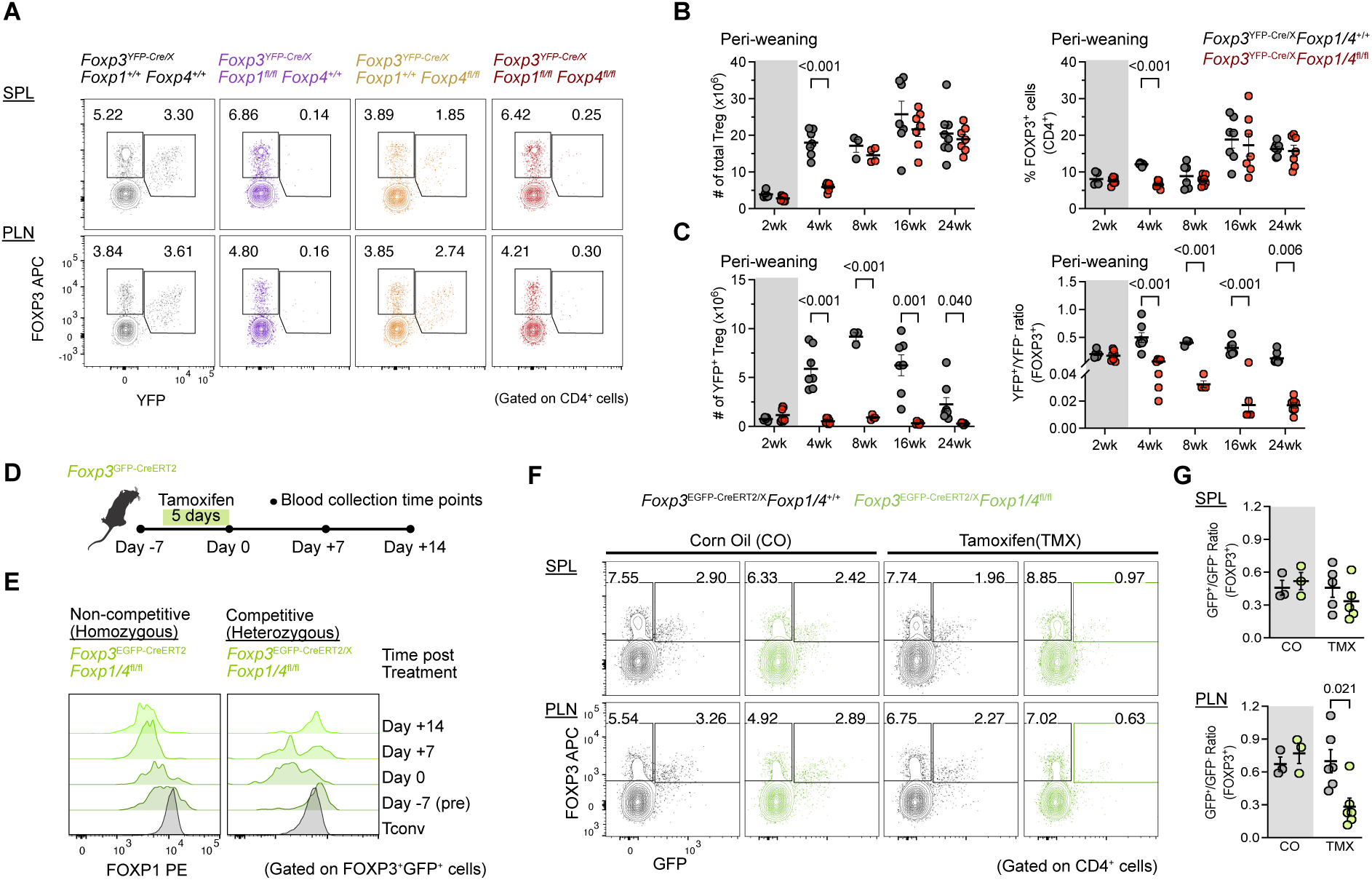
Loss of FOXP1 or both FOXP1 and FOXP4 in Treg cells leads to a competitive disadvantage. (A) Representative flow cytometric plots of the percentage of Treg cells in gated CD4^+^ cells from spleen (SPL) and peripheral lymph nodes (PLN), collected from females heterozygous of all four strains at 8 weeks of age. (B and C) Longitudinal analysis of cellularity and frequencies of splenic FOXP3^+^ Treg within CD4^+^ T cells (B) and the YFP^+^/YFP^−^ ratio (C) at the indicated time points. Each dot represents one mouse (n= 5 to 8). (D) Schematic plot of workflow and tamoxifen-induced acute FOXP1/FOXP4 depletion and blood collection time points. (E) Representative flow cytometric analysis showing dynamic changes in the proportion of FOXP3^+^GFP^+^FOXP1^lo/–^ cells in peripheral blood before and after tamoxifen-induced acute depletion in *Foxp3^EGFP-^*^CreERT2^ mice. (F) Flow cytometry plots showing the frequencies of FOXP3^+^YFP^+^ Treg cells within CD4^+^ cells, collected from *Foxp3^EGFP-^*^CreERT2/X^ heterozygous and age-matched controls on day 7 post-tamoxifen treatment. (G) Quantification of the GFP^+^/GFP^−^ ratio in gated CD4^+^ cells from (F) (n= 3 to 6). Data are presented as mean ± SEM; each dot represents data from an individual mouse. Statistical analysis: two tailed unpaired *t*-test (B, C and G). *P* values are shown for significant difference (*P* < 0.05).

To ensure that the competitive disadvantage of FOXP1/FOXP4 deficient Tregs was not a consequence of weaning or thymic developmental differences, we used a temporally inducible knockout model to acutely delete *Foxp1* and *Foxp4* in mature, peripheral Tregs. Tamoxifen was administered to mice carrying the EGFP-Cre-ERT2 fusion cassette at the 3’-UTR of *Foxp3,* which allowed us to acutely deplete FOXP1 and FOXP4 in FOXP3-expressing cells in adult mice (Fig. 1D). We identified deleted cells by flow cytometry based on the expression of EGFP and the loss of FOXP1 protein. When CreERT2 is expressed on both X chromosomes and all Tregs undergo deletion, FOXP1/FOXP4-deficient Tregs (FOXP3^+^GFP^+^FOXP1^lo/–^) persist in peripheral blood until at least day 14 post-tamoxifen (Fig. 1E). In contrast, when the CreERT2 is heterozygous approximately half of the cells retain FOXP1/FOXP4 expression after tamoxifen dosing setting up a direct competition for needed growth/survival factors. In the competitive environment, the presence of FOXP1^lo/–^ Treg cells in the blood was only transient. And by day 14, circulating FOXP1-sufficient Tregs had completely repopulated suggesting that lack of FOXP1 and FOXP4 results in a competitive disadvantage within the Treg pool (Fig. 1E). Similar to the peripheral blood, the competitive disadvantage seen with loss of FOXP1 and FOXP4 was observed in SLOs (Fig. 1, F and G).

### FOXP1/FOXP4 are required for peripheral Treg survival

To further investigate the basis of the low numbers and proportion of FOXP1/FOXP4 deficient Tregs in a competitive environment, we assessed processes of entry or exit from this pool. Tregs can enter the circulating pool either through development in the thymus (tTregs) or through conversion of peripheral naïve CD4 T cells (pTregs) and exit through loss of FOXP3 to become exTregs or cell death. The decreased proportion of FOXP1/FOXP4-deficient Treg cells was not due to decreased thymic development. While there was a transient decrease in YFP^+^ plus YFP^-^FOXP3^+^ thymic Tregs at 4 weeks of age in the absence of FOXP1/FOXP4, there was no statistical difference in the relative proportion of deleted cells (fig. S2A). Furthermore, there was no statistical difference of YFP^+^ relative to YFP^-^ in Helios^+^Nrp1^+^ tTreg isolated from SLOs (fig. S2, B and C) suggesting normal tTreg seeding of the periphery. To directly assess input from the pTreg compartment we used a mixed, adoptive transfer approach that allowed us to track donor derived pTreg at steady state. We sorted naïve T (Tnv) cells (CD4^+^CD25^-^YFP^-^CD44^lo^CD62L^hi^) from either *Foxp3^YFP-^*^Cre/Cre^*Foxp1/4*^+/+^ or *Foxp3*^YFP*-*Cre/Cre^*Foxp1/4*^fl/fl^ mice (CD45.2, CD90.2) and mixed them at 1:1 ratio with Tnv cells from congenic *Foxp3*^YFP*-*Cre/Cre^*Foxp1/4*^+/+^ mice (CD45.1, CD90.2), followed by adoptive transfer into congenically disparate *Foxp3*^EGFP-CreERT2^ (CD45.2, CD90.1) recipients (Fig. 2A). At 6 weeks after the transfer, approximately 0.5-2% of donor-derived FOXP1/FOXP4 sufficient cells were pTregs based on YFP expression (Fig. 2B). In contrast, *Foxp3*^YFP-Cre/Cre^*Foxp1/4*^fl/fl^-derived (CD45.2^+^) pTregs were nearly undetectable (Fig. 2, B and C). These data do not differentiate between the decreased pTreg seen at 6 weeks post-transfer being a consequence of a failure to generate the pTreg lacking FOXP1 and FOXP4 or an inability to maintain them over the time frame of the experiment. To distinguish between these possibilities, we mimicked the pTreg generation process by co-culturing a 1:1 mixture of congenically disparate *Foxp3*^YFP*-*Cre/Cre^*Foxp1/4*^+/+^ and *Foxp3*^YFP*-*Cre/Cre^*Foxp1/4*^fl/fl^ Tnv cells under iTreg-polarizing conditions for 72 hours (fig. S3A). FOXP3 was detectable in approximately 25% and 70% of cells after 24 and 48 hours of culture, respectively (fig. S3B). By 72 hours after stimulation, iTregs from *Foxp3*^YFP*-*Cre/Cre^*Foxp1/4*^fl/fl^ mice exhibited a noticeable competitive disadvantage (fig. S3, C and D). The kinetics of the loss of *Foxp3*^YFP*-*Cre/Cre^*Foxp1/4*^fl/fl^ relative to control cells corresponded with decreased FOXP1 and occurred despite similar rates of division based on dye dilution (fig. S3E). Taken together, these findings suggest the observed competitive disadvantage in *Foxp3*-driven pTreg lacking FOXP1 and FOXP4 is due to a lack of persistence rather than differences in thymic or peripheral generation.

**Fig. 2.**
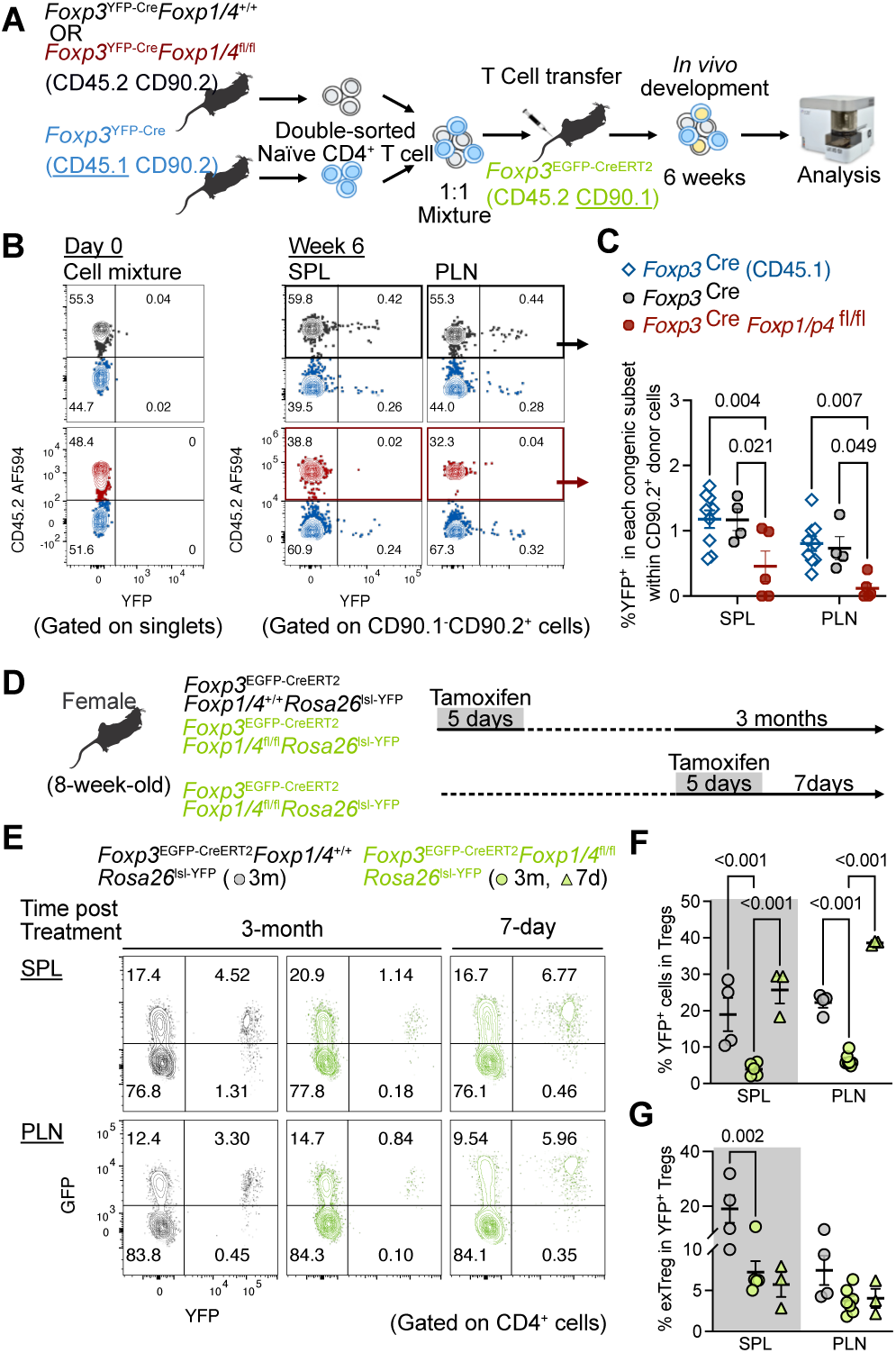
Defective peripheral Treg maintenance in mice lacking FOXP1 and FOXP4. (A) Schematic workflow for investigating *in vivo* pTreg competitiveness at steady state. Naïve CD4^+^ from *Foxp3*^YFP-Cre^*Foxp1/4*^+/+^ and *Foxp3*^YFP-Cre^*Foxp1/4*^fl/fl^ were isolated, mixed at a 1:1 ratio with congenically disparate *Foxp3*^YFP-Cre^ cells and adoptively transferred into *Foxp3*^EGFP-CreERT2^ mice. Cells from spleen and peripheral LN were analyzed 6 weeks later. (B) Representative flow cytometric plots assessing the proportion of YFP^+^ pTreg in CD90.2^+^ donor cells before and 6 weeks after adoptive transfer. (C) Quantification of the percentage of YFP^+^ pTreg in each donor compartment 6 weeks after adoptive transfer (n=4 to 9). (D) Schematic workflow for assessing Treg persistence in the periphery. Mice were treated for 5 days with tamoxifen then assessed 7 days or 3 months after deletion. (E) Representative flow cytometric plots of the frequency of peripheral persistent YFP^+^ Treg generated during the tamoxifen pulse. (F) Quantification of persistent YFP^+^ Treg and (G) percentage of Treg cells that convert into exTreg (n=3 to 7). Data are presented as mean ± SEM, each dot represents data from an individual mouse. Statistical analysis: 2-way ANOVA. *P* values are shown where p<0.05.

To assess the *in vivo* persistence and stability of Tregs, we crossed the *Rosa26*^lsl-YFP^ fate-mapper into *Foxp3^EGFP-Cre-ERT2^* and *Foxp3^EGFP-Cre-ERT2^Foxp1/4*^fl/fl^ mice. In *Foxp3^EGFP-Cre-ERT2^Foxp1/4*^fl/fl^*Rosa26*^lsl-YFP^ mice treated with tamoxifen, the EGFP-Cre-ERT2 fusion protein translocates into the nucleus to delete *loxP*-flanked *Foxp1*, *Foxp4,* and the transcriptional *STOP* that blocks YFP expression within the same GFP^+^ Treg (fig. S4, A and B). That allowed us to track the stability of YFP^+^ FOXP1/FOXP4-deficient Tregs generated during the tamoxifen treatment period. Age-matched females were treated with tamoxifen for 5 consecutive days, and SLOs were collected and analyzed 1 week and 3 months later (Fig. 2D). Seven days after tamoxifen, equivalent percentages of GFP^+^ (FOXP3^+^) Treg from both FOXP1/FOXP4-sufficient and FOXP1/FOXP4-deficient mice expressed YFP (fig. S4A). However, by 3 months post-tamoxifen, there were substantially fewer YFP^+^GFP^+^ Treg derived from *Foxp3*^EGFP*-*Cre-ERT2^*Foxp1/4*^fl/fl^ compared with controls in spleen and PLNs (Fig. 2, E and F). Furthermore, FOXP1/FOXP4-deficient Tregs and wild-type Tregs had comparable levels of conversion into YFP^+^GFP^-^ exTregs in these tissues (Fig. 2, E and G).

The decreased persistence of FOXP1/FOXP4 deficient Treg may be due to decreased homeostatic proliferation or increased cell death. *In vitro* iTreg dye-dilution (fig. S3E), *in vivo* BrdU incorporation (Fig. 3, A and B) and measurement of Ki67 expression (Fig. 3C) showed little if any difference between FOXP1/FOXP4-sufficient and FOXP1/FOXP4-deficient Treg, indicating intact rates of homeostatic proliferation. In contrast, YFP^+^ Treg from *Foxp3*^YFP*-*Cre/X^*Foxp1/4*^fl/fl^ had substantially more late-stage apoptotic and necrotic cells compared with those from control animals (Fig. 3, D to F). Consistent with increased cell death, Caspase 3/7 activity was elevated in splenic *Foxp3*^YFP*-*Cre/X^*Foxp1/4*^fl/fl^ YFP^+^ Treg. (Fig. 3G). Together these data indicate: (1) The absence of FOXP1/FOXP4 leads to a loss of competitive fitness at steady state; (2) FOXP1/FOXP4 are dispensable for Treg development but required for maintaining Tregs in the peripheral T cell pool; (3) decreased survival of FOXP1/FOXP4-deficient Tregs is the primary mechanism underlying their competitive disadvantage in SLOs.

**Fig. 3.**
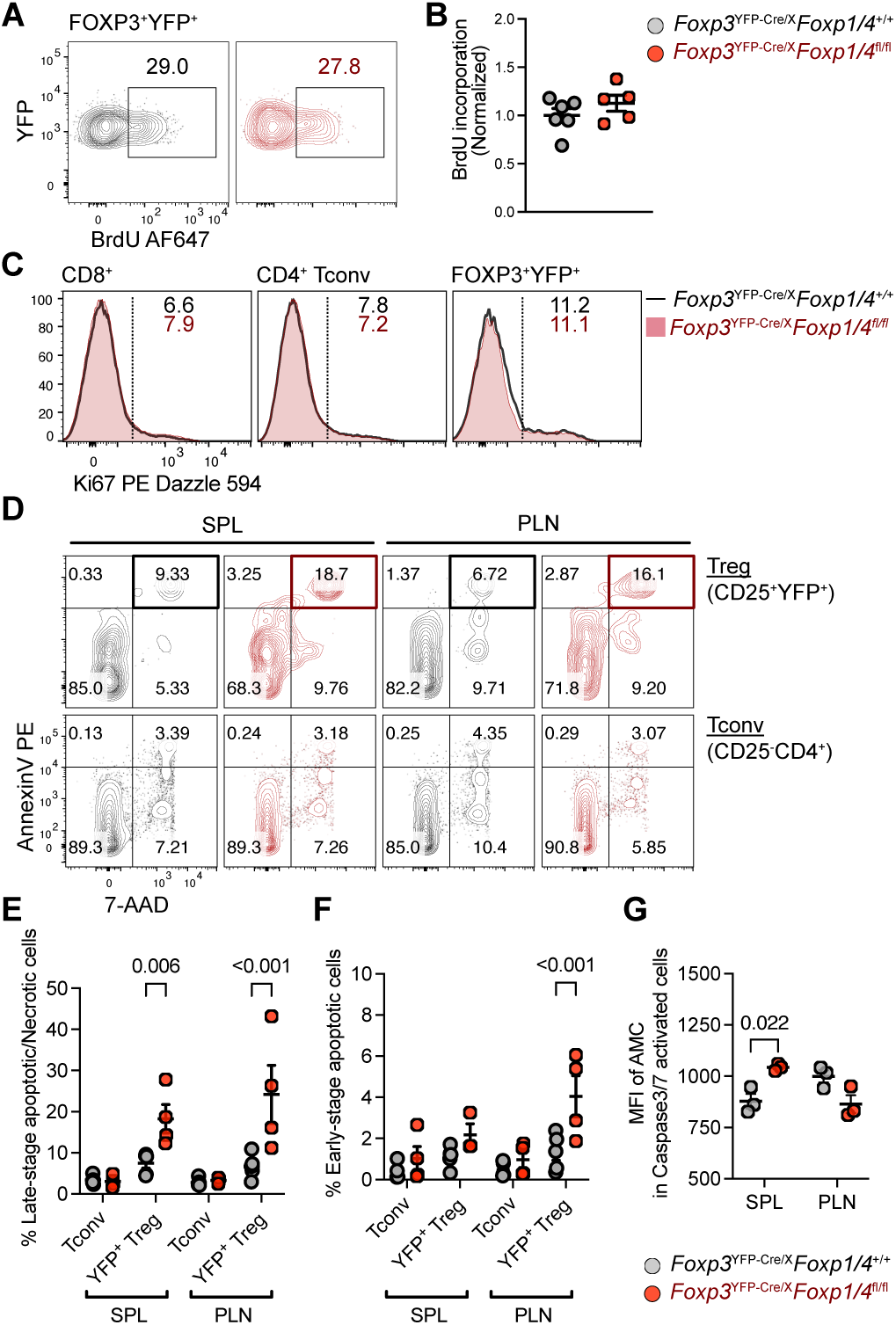
Reduced viability of FOXP1/FOXP4-deficient Tregs. (A) Representative flow cytometric plots of BrdU incorporation in YFP⁺ Tregs and (B) quantification (n = 5 to 6 per group). (C) Histogram of percentage of Ki67^+^ cells in *Foxp3*^YFP-Cre/X^ (black line) and *Foxp3*^YFP-Cre/X^ *Foxp1/Foxp4*^fl/fl^ (red shade), representative of three independent experiments. (D) Flow cytometric plots of cell viability in YFP^+^ Treg and Tconv. Data are representative of three independent experiments. (E and F) Quantification of the frequency of (E) late-stage apoptotic/necrotic cells (F) and early-stage apoptotic cells (n=4). (G) Quantification of caspase 3/7 activity in YFP^+^ Treg (n=3 to 4). Data are presented as mean ± SEM, each dot represents data from an individual mouse. Statistical analysis: two tailed unpaired *t* -test. *P* values are shown for significant difference.

### Gene expression and chromatin accessibility changes in FOXP1/FOXP4-deficient Tregs

We next sought to determine the genetic changes that led to reduced survival of FOXP1/FOXP4-deficient Tregs. Bulk RNA-seq of sorted YFP^+^ Treg from *Foxp3*^YFP*-*Cre/X^*Foxp1/4*^+/+^ and *Foxp3*^YFP*-*Cre/X^*Foxp1/4*^fl/fl^ mice identified 1210 differentially expressed genes (DEGs) (Fig. 4A and table S1). Gene Ontology and KEGG enrichment analysis indicated that many DEGs were involved in cytokine binding, immune receptor activity, and protein tyrosine kinase activity (fig. S5, A to C). Given the survival defect, we separately evaluated gene expression related to programmed cell death pathways and identified altered expression of several genes involved in apoptosis, necroptosis, ferroptosis and autophagy in Tregs from *Foxp3*^YFP*-*Cre/X^*Foxp1/4*^fl/fl^ compared with controls. With the exception of *Fth1,* these alterations did not reach statistical significance (fig. S5D).

**Fig. 4.**
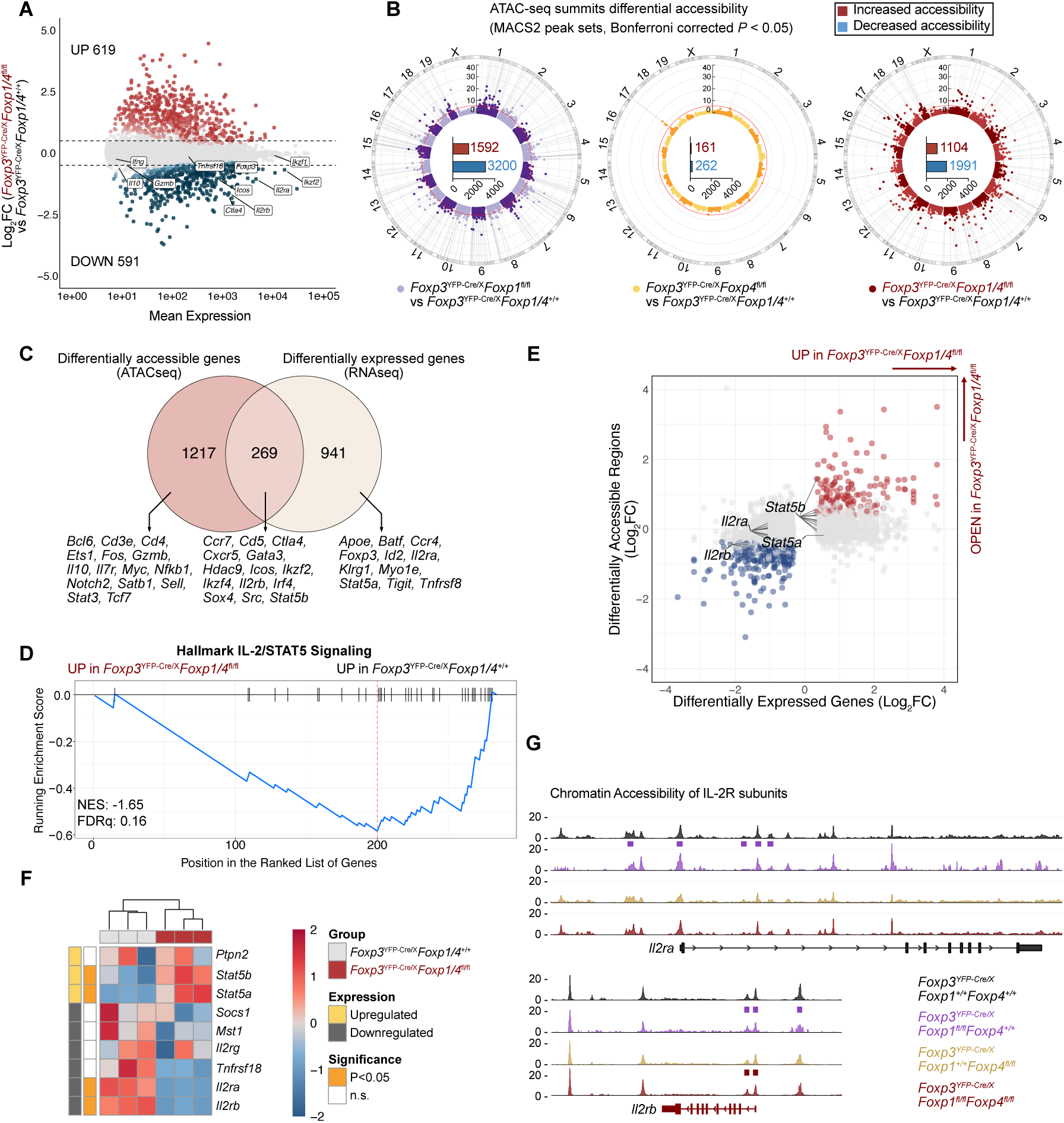
Chromatin accessibility and transcriptome analysis reveal defective IL-2/STAT5 signaling in FOXP1/FOXP4-deficient Treg cells. (A) Shrunken MA-plot comparing the gene expression in YFP^+^ Tregs sorted from *Foxp3*^YFP-Cre/X^*Foxp1/4*^+/+^ and *Foxp3*^YFP-Cre/X^*Foxp1/4*^fl/fl^. DEGs are shown in red (Up) and blue (Down). The exact number of DEGs and gene symbols are labelled. (B) Circle Manhattan plot showing the landscape of chromatin accessibility at the whole genome level. Data are represented as relative ratio: *Foxp3*^YFP-Cre/X^*Foxp1*^fl/fl^ vs *Foxp3*^YFP-Cre/X^ (purple dots); *Foxp3*^YFP-Cre/X^*Foxp4*^fl/fl^ vs *Foxp3*^YFP-Cre/X^ (yellow dots); and *Foxp3*^YFP-Cre/X^*Foxp1/Foxp4*^fl/fl^ vs *Foxp3*^YFP-Cre/X^ (red dots). Radial scale represents -log_10_ adjusted *P*-values and the bar plot in the center represents the number of peaks with adjusted-*P*<0.05. (C) Venn diagram of differentially accessible regions and differentially expressed genes from Treg cells. The number is shown as the unique gene symbols. Selected gene loci are listed below the diagram. (D) Ranked enrichment plot of IL-2/STAT5 signaling in gene set enrichment analysis of shared genes. (E) Scatter plot of 1210 DEGs with chromatin accessibility with selected genes of interest indicated (red indicates increased chromatin accessibility and transcription; blue indicates decreased accessibility and transcription; gray represents others). (F) Heatmap of transcription level of IL-2/STAT5 signaling-related genes. (G) ATAC-seq signal peaks at the *Il2ra* and *Il2rb* loci in Treg cells from all four strains. Peaks that show significant difference compared with wild-type Tregs are indicated with a box above. Two biological replicates were used for each genotype.

We also assessed cell-autonomous chromatin accessibility in Tregs that were deficient in FOXP1 and/or FOXP4. Approximately twice as many loci exhibit decreased compared with increased accessibility when FOXP1, FOXP4 or both were absent; the most changes were evident with the isolated absence of FOXP1 (Fig. 4B and table S2). Comparing RNA-seq and ATAC-seq data from *Foxp3*^YFP*-*Cre/X^*Foxp1/4*^fl/fl^ with wild-type Tregs, we identified 269 genes (22.2% of total DEG) with differences in both gene expression and chromatin accessibility (Fig. 4C), including *Ccr7, Ctla4, Gata3, Icos,* and *Ikzf2*, which are key regulators of Treg phenotype and function. Gene set enrichment analysis (GSEA) of these 269 genes against Hallmark pathways identified only the IL-2/STAT5 signaling pathway as significantly enriched (Fig. 4D).

To identify potential direct targets of FOXP1/FOXP4 transcriptional regulation, we focused on those loci with discordant gene expression and chromatin accessibility changes (i.e., those with altered steady state RNA levels but without a change in chromatin accessibility, those where chromatin was open but gene expression was decreased and those where chromatin was more closed with increased gene expression) (Fig. 4E, loci noted in grey). We then focused our analysis on genes included in the IL-2/STAT5 signaling module (Fig. 4F). The *Il2ra* locus is an example of direct transcriptional regulation; *Il2ra* mRNA was significantly downregulated in *Foxp3*^YFP*-*Cre/X^*Foxp1/4*^fl/fl^ Tregs (Fig. 4F) but its chromatin accessibility did not differ significantly from the control (Fig. 4G). In contrast, the *Il2rb* locus exhibited reduced chromatin accessibility at both its promoter and upstream regions in *Foxp3*^YFP*-*Cre/X^*Foxp1/4*^fl/fl^ and *Foxp3*^YFP*-*Cre/X^*Foxp1*^fl/fl^ Tregs and decreased transcript levels, suggesting epigenetic rather than transcriptional regulation (Fig. 4G). These data suggest that the mechanism underlying decreased gene expression resulting from deletion of *Foxp1* on *Il2ra* is transcriptional while its effect on *Il2rb* expression is epigenetic and suggested defects in IL-2/STAT5 related signaling pathways.

### Impaired IL-2/STAT5 signaling in FOXP1/FOXP4-deficient Tregs

We next assessed the expression of each subunit of the high-affinity IL-2Rαβγ in YFP⁺ Tregs. The expression level of IL-2Rα (CD25) was markedly reduced in YFP^+^ Treg from *Foxp3*^YFP*-*Cre/X^*Foxp1/4*^fl/fl^ and *Foxp3^YFP-^*^Cre/X^*Foxp1*^fl/fl^ but not *Foxp3^YFP-^*^Cre/X^*Foxp4*^fl/fl^ mice (Fig. 5A and fig. S6A). Cell surface levels of IL-2Rβ (CD122) and IL-2Rγ (CD132) were slightly higher in Treg cells lacking FOXP1/FOXP4 suggesting intact expression of the intermediate affinity form of the IL-2R (Fig. 5B). Consistent with decreased expression of the high affinity IL-2Rαβγ and intact expression of the intermediate affinity IL-2Rβγ, fewer FOXP1/FOXP4-deficient Tregs exhibited responsiveness to low-dose IL-2 (0.4 IU/mL) as assessed by STAT5 phosphorylation; this was largely overcome with high-dose IL-2 (40 IU/mL) (Fig. 5C and 5D). In those Tregs with phosphorylated STAT5, the level of pSTAT5 in response to either dose of IL-2 was comparable regardless of FOXP1/FOXP4 expression (Fig. 5C and 5E). Acute depletion of FOXP1/FOXP4 in mature Tregs similarly resulted in decreased CD25 expression and fewer cells phosphorylating STAT5 in response to low dose, but not high dose IL-2 (fig. S6, B to E). Together, these observations support a model in which the primary challenge to survival in FOXP1/FOXP4-deficient Tregs is a decrease in surface CD25, with a resulting decline of IL-2 responsiveness.

**Fig. 5.**
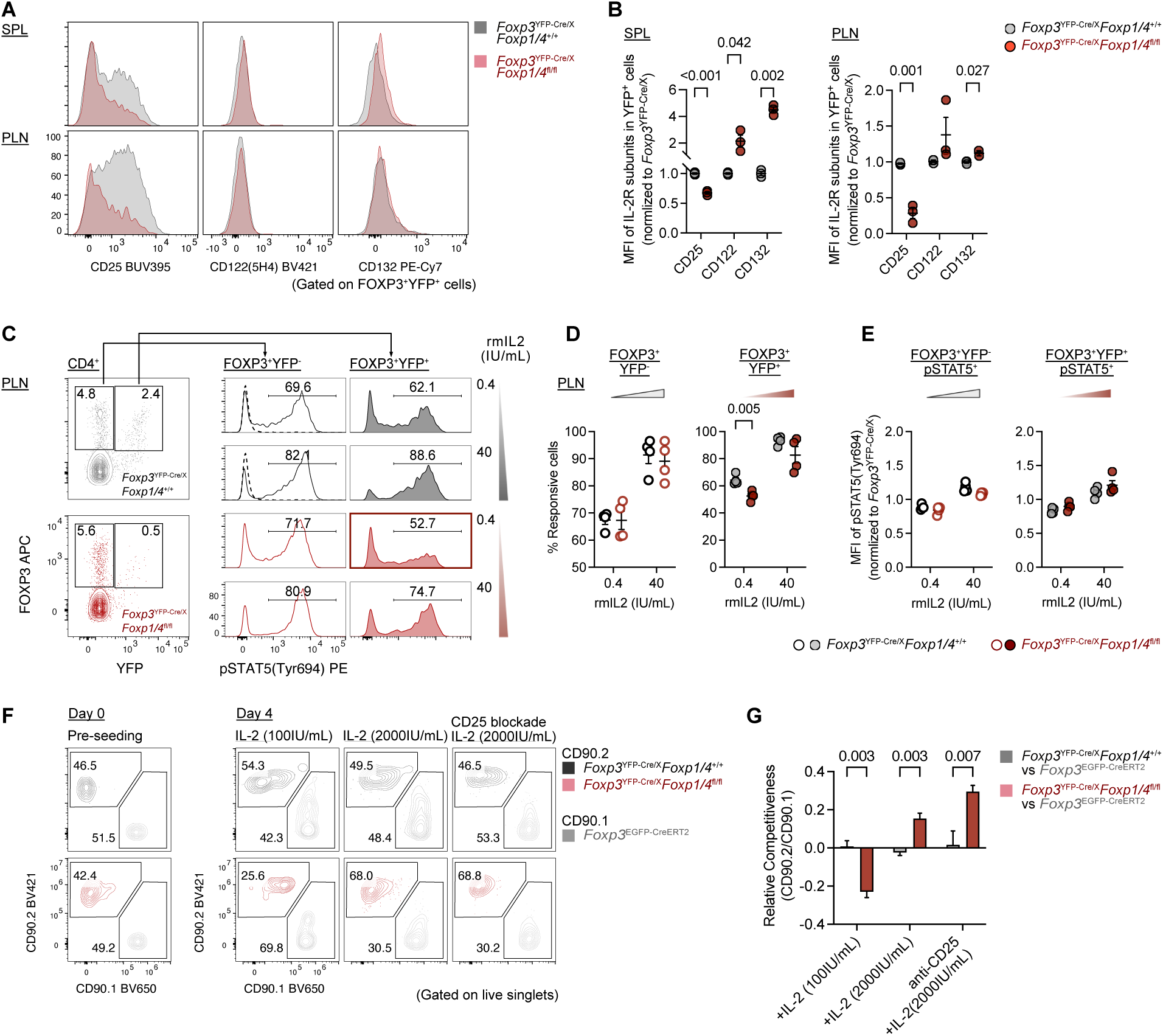
Deletion of FOXP1/FOXP4 reduces IL-2 responsiveness of Tregs. (A) Histograms comparing the cell surface expression levels of IL-2R subunits in FOXP3⁺YFP⁺ gated cells from *Foxp3*^YFP-Cre/X^ (gray shaded) and *Foxp3*^YFP-Cre/X^*Foxp1/Foxp4*^fl/fl^ (red shaded) mice. Data are representative of three independent experiments. (B) Relative expression level of IL-2R subunits on the surface of YFP⁺ cells from the spleen and PLN (n = 3 per group). Data are representative of three independent experiments. (C) Representative flow cytometric plots showing pSTAT5 levels in PLN Tregs in response to rmIL-2 stimulation at low and high doses. (D) Quantification of the proportion of IL-2-responsive cells and (E) relative pSTAT5 levels in Tregs from (C) (n = 4 per group). (F) Representative flow cytometric plots showing the relative proportion of donor cells after 4 days of co-culture with treatments. (G) Calculated relative competitiveness with different treatments. Data are presented as mean ± SEM, each dot represents data from an individual mouse. Statistical analysis: two tailed unpaired *t* -test (B, D, E and G). *P* values are shown for significant difference.

To confirm that the impaired IL-2 response directly contributed to the competitive disadvantage of FOXP1/FOXP4-deficient Tregs, we co-transferred *ex vivo* expanded YFP^+^ nTregs from *Foxp3*^YFP-Cre/X^*Foxp1/4*^+/+^ or *Foxp3*^YFP-Cre/X^*Foxp1/4*^fl/fl^ (CD45.2, CD90.2) mixed with congenic *Foxp3*^EGFP-CreERT2/X^*Foxp1/4*^fl/fl^ (CD45.2, CD90.1) into CD45.1 *Tcrb/Tcrd*^-/-^ hosts, which lack an endogenous source of IL-2(*17*) (fig. S7A). In this IL-2-deficient setting, the competitive disadvantage of FOXP1/FOXP4-deficient Treg was no longer observed (fig. S7B). To determine if providing supraphysiological IL-2 could rescue the competitive disadvantage of FOXP1/FOXP4-deficient Tregs, we co-cultured FOXP1/FOXP4-sufficient and deficient Tregs with relatively low (100 IU/mL) versus high (2000 IU/mL) dose IL-2. In contrast to the competitive disadvantage seen with low dose IL-2, FOXP1/FOXP4-deficient Treg outcompeted control cells with the higher dose of IL-2 (Fig. 5F and G), potentially due to increased cell surface expression of the IL2-Rβγ. This was not inhibited by anti-CD25 treatment suggesting a significant contribution of the intermediate affinity IL-2Rβγ. Collectively, these findings demonstrate that reduced IL-2 responsiveness due to diminished CD25 expression is the primary cause of the impaired fitness of FOXP1/FOXP4-deficient Tregs.

### FOXP1 is necessary for *Il2ra* super-enhancer activity via chromatin looping

We provide evidence above that FOXP1 and FOXP4 are necessary for maintenance of high levels of cell-surface CD25 (Fig. 5A and S6B). In contrast, iTregs generated under competitive conditions from *Foxp3*^YFP-Cre^*Foxp1/4*^fl/fl^ Tnv were capable of wild-type levels of CD25 induction during the first 48 hours post-polarization (Fig. 6A). To confirm this was not a consequence of deletion occurring after stimulation of naïve cells, we generated FOXP1/FOXP4-deficient Tnv from tamoxifen-treated *Cd4*^CreERT2^*Foxp1/4^f/f^* mice (fig. S8) and subjected them to iTreg polarization. After 48 hours of iTreg induction, FOXP1/FOXP4-deficient *Cd4*^CreERT2^ iTreg cells also showed wild-type levels of CD25 and a trend towards higher CD25 levels/cell (Fig. 6B and 6C). This suggested that expression of CD25 on Tregs is initially FOXP1/FOXP4-independent but that the maintenance of high expression levels is dependent on FOXP-factors.

**Fig. 6.**
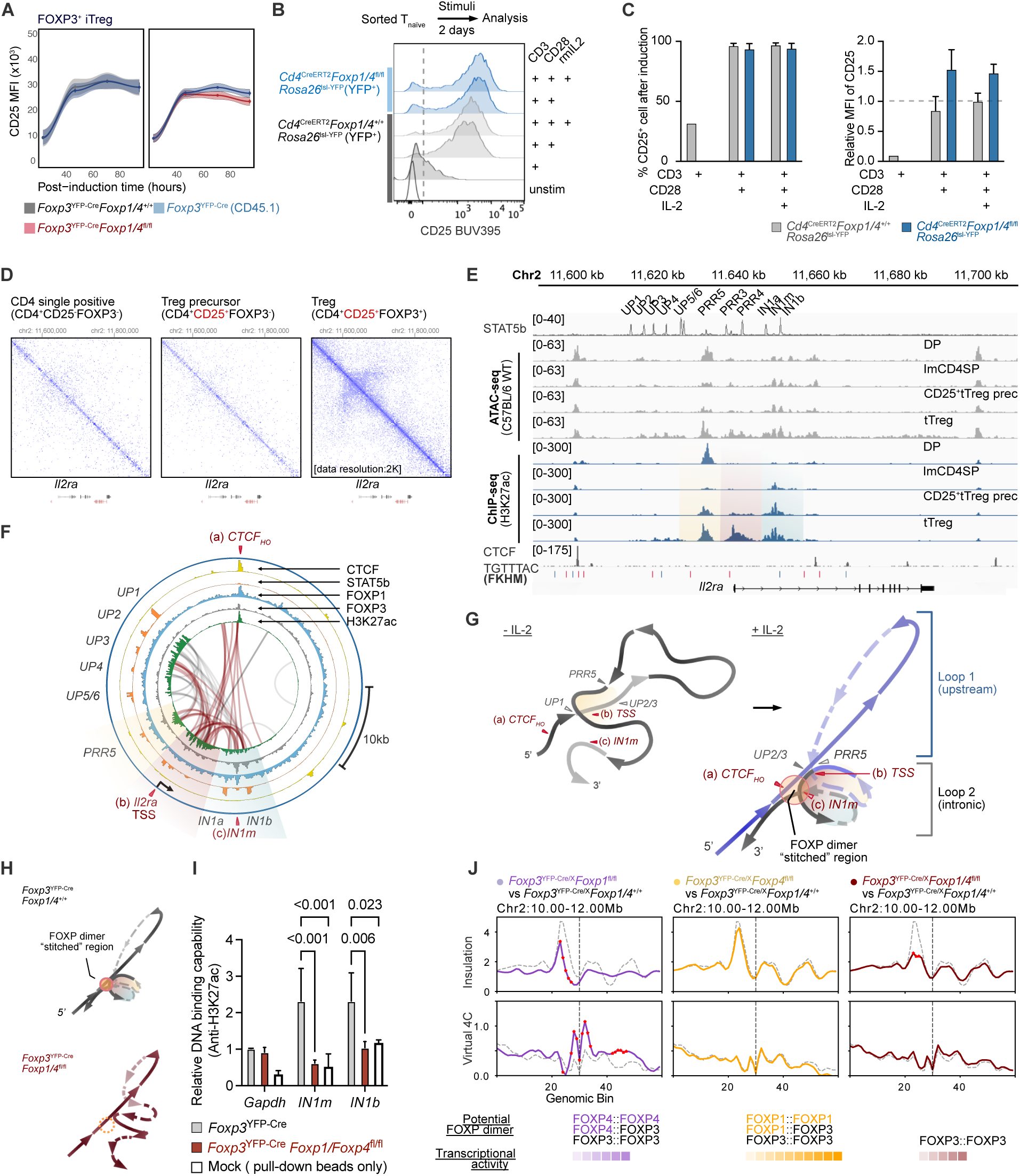
FOXP1-associated chromatin architecture is essential for maintaining *Il2ra* transcription in mature Tregs. (A) Expression level of CD25 in FOXP3-expressing cells during the iTreg polarization under competitive conditions. Curves represent locally estimated scatterplot smoothing for each condition, with points and error bars indicating mean ± SEM at each time point. The genotypes of cell origins are indicated by colors. (B) Histogram of the relative MFI of CD25 in iTregs polarized from FOXP1/FOXP4-deficient naïve T cells with various stimuli. Data are representative of two independent experiments. (C) Quantification of the percentage (left) and relative expression level of CD25 in (B). MFI of CD25 was normalized to *Cd4*^CreERT2^*Foxp1/p4*^+/+^*Rosa26*^lsl-YFP^ Tnv cells stimulated with CD3/CD28/IL-2. (D) In-situ Hi-C plots at the *Il2ra* locus during thymic Treg development (samples GSM6705657, GSM6705655, and GSM6705675). (E) ATAC-seq and H3K27ac ChIP-seq tracks at the *Il2ra* locus during Treg development (samples GSM2734684, SRR5385310, SRR5385309, SRR5385308, SRR5385307, SRR5385355, SRR5385353, SRR5385351, SRR5385349, and GSM7213946). STAT5B binding sites are denoted in the top track. Forkhead consensus motif (FKHM) sequences are denoted below the tracks. (F) RNA-PolII ChIA-PET plot illustrating chromatin interactions (bridging loops, newly established interactions after IL-2 stimulation are shown in red) within the *Il2ra* locus in wild-type CD4 cells. The plot includes overlaid ChIP-seq peaks obtained in Tregs. Shaded yellow, pink and blue areas correspond to similar shading in (E). (G) Schematic illustration of FOXP-instructed chromatin folding, with and without IL-2 stimulation, into a topologically associating chromatin structure. Color-shaded areas indicate corresponding H3K27ac-enriched transcriptionally active regions as shown in e and f. Critical FOXP-independent (grey triangle) and FOXP-associated (red triangle) FKHM regions marked. (H) Hypothesized chromatin interaction loss at *Il2ra* in the absence of FOXP1/FOXP4. (I) Quantified DNA-binding at *IN1m* and *IN1b* by ChIP-qPCR in FOXP1/FOXP4 expressing (grey) and FOXP1/FOXP4 deficient (red) Foxp3^+^ Tregs. (J) Calculated insulation score (top) and virtual 4C (bottom) derived from Treg-specific trained model predicted interaction matrices for *Foxp3*^YFP-Cre/X^*Foxp1*^fl/fl^, *Foxp3*^YFP-Cre/X^*Foxp4*^fl/fl^ and *Foxp3*^YFP-Cre/X^*Foxp1/Foxp4*^fl/fl^ mice at Chr2:11,410,143-11,878,893 (mm10), shown as difference from *Foxp3*^YFP-Cre/X^*Foxp1/Foxp4*^+/+^ control (dotted line). The vertical lines indicate the *Il2ra* TSS, and red asterisks denote statistically significant difference compared to control. Data are presented as mean ± SEM. Statistical analysis: two tailed unpaired *t* -test (C), 2-way ANOVA with Dunnett’s post-hoc test (I). *P* values are shown for significant difference.

The experiments above demonstrate that maintenance of *Il2ra* transcription and subsequent CD25 surface expression are dependent on continued FOXP1 and FOXP4 expression and suggest this is a direct transcriptional mechanism. Recent studies have pointed to the *Il2ra* super-enhancer and its ability to form enhancer-promoter loops(*18–20*). Re-analysis of a publicly accessible in-situ Hi-C dataset (table S3) identified a topologically associating domain (TAD) structure in the *Il2ra* promoter-intronic region of CD4^+^CD25^+^FOXP3^+^ mature thymic Tregs that was absent in both CD4^+^CD25^−^FOXP3^−^ single positives and CD4^+^CD25^+^FOXP3^-^ Treg precursors (Fig. 6D).

Increasing chromatin accessibility and H3K27 acetylation within the *Il2ra* intronic region correlated with the TAD buildup during Treg development (Fig. 6E). Within this intronic region, reanalysis of publicly available ChIP-seq data revealed that both FOXP1 and FOXP3 were found at the STAT5 *IN1m* and *IN1b* response elements in Treg cells (Fig. 6F and fig. S9A). Additionally, this intronic region exhibited an active enhancer marked by binding of NFAT1 and the chromatin remodeler p300(*21*) in addition to acetylated H3K27 (fig. S9A). Motif analysis further revealed that conventional forkhead binding motifs (FKHM, TGTTTAC) are enriched between *Il2ra*-locus flanking CTCF sites (11,600-11,670kb, bottom of both Fig. 6E and fig. S9A), suggesting a potential role of FOXP proteins in bridging distant DNA together to form a TAD-associated super-enhancer.

Combining our previously reported RNA polymerase-II (RNA Pol II)-ChIA-PET analysis(*18*) in CD4^+^ T cells with the ChIP-seq data from Tregs suggested three potential FOXP anchor points: the high-occupancy CTCF site (CTCF_HO_, sites that are high-affinity and stably bound by CTCF(*22*)), the transcriptional start site (TSS), and the intronic IN1m region (Fig. 6F). The CTCF_HO_ to TSS and CTCF_HO_ to the intronic region loops are not present in the absence of an IL-2 stimulus (fig. S9B). We hypothesized that a topologically associated chromatin organized complex involving these sequences might be critical for persistent *Il2ra* transcription (Fig. 6G) and function in the following manner: During TAD formation in response to IL-2, FOXP1 and/or FOXP3 participate in the assembly of a protein complex bridging IN1m and CTCF_HO_ to form the required interactions. This spatial network provides a structural foundation for the super-enhancer complex that includes STAT5, NFAT1, and other co-activators. The lack of STAT5 bound to either intronic IN1a/IN1b sites in Tregs(*20*) or to the CTCF sites in CD4 cells(*23*) leads to a disrupted TAD and a decline in CD25 expression. Based on this model, we further hypothesized that the absence of FOXP1/FOXP4 would lead to in an inability to form the required looping, resulting in decreased enhancer activity within the intronic region (Fig. 6H). To test this hypothesis, we performed ChIP-qPCR for H3K27 targeting the *IN1m* and *IN1b* regions in YFP^+^ cells from *Foxp3*^YFP*-*Cre^*Foxp1/4*^+/+^ and *Foxp3*^YFP*-*Cre^*Foxp1/4*^fl/fl^ mice. Reduced enrichment of the active enhancer (H3K27ac) marks at these intronic regions in FOXP1/FOXP4-deficient Tregs (Fig. 6I) supports a model of *Il2ra* transcriptional regulation anchored by FOXP proteins at these sites. Analysis of Tconv cells and “Treg wannabe” cells(*4*), two populations of T cell lacking FOXP3, demonstrated that the presence of FOXP3 was not necessary for formation of the *Il2ra* locus TAD (fig. S9C). Lastly, to determine whether the enhancer activity is chromatin conformation associated, we performed *in silico* Hi-C analysis using a model trained in mouse Tregs (fig. S10, A and B). Using the predicted chromatin interaction matrix and calculated insulation scores, there was a reduction of TAD in regions adjacent to *Il2ra* TSS in both *Foxp3*^YFP*-*Cre/X^*Foxp1*^fl/fl^ and *Foxp3*^YFP*-*Cre/X^*Foxp1/4*^fl/fl^ Tregs, while *Foxp3*^YFP*-*Cre/X^*Foxp4*^fl/fl^ is similar to wild-type control (Fig. 6J).

## DISCUSSION

Tregs hold considerable promise for therapeutic applications, yet details of the molecular requirements for their persistence and competitiveness have not been fully explored. In this study, we investigated the cell-intrinsic role of FOXP1/FOXP4 under homeostasis and show that: (1) FOXP1/FOXP4 are required to maintain Tregs within the peripheral T cell pool, (2) the competitive disadvantage observed in FOXP1/FOXP4-deficient Treg cells is mainly ascribed to the loss of FOXP1, (3) reduced CD25 expression limits responsiveness to IL-2 in FOXP1/FOXP4-deficient Tregs, (4) *Il2ra* transcription has two distinct stages and FOXP1 is essential for maintaining high levels of CD25 expression, and (5) FOXP1 contributes to a chromatin looping structure, facilitating pSTAT5-related enhancer activity within the *Il2ra* intronic region.

Treg development relies on coordinated TCR and IL-2 signaling. Using the *Tokey* timer reporter system, David and colleagues demonstrated that the majority of thymic Tregs arise from FOXP3⁻CD25^hi^ precursors, which are the earliest lineage subset to receive TCR stimulation, and effects that persist throughout the process of Treg lineage commitment(*24*). Intrathymic transfer experiments further showed that FOXP3⁻CD25^hi^ precursors that are poised to upregulate FOXP3 expression require only IL-2 or IL-15 signals without further TCR engagement(*25*). Additionally, exogenous IL-2 or selective activation of STAT5 in H-2DMa or T-cell specific SLP-76 knockout mice can partially restore Treg development independent of TCR signaling(*26*). Moreover, FOXP3 itself has been reported to physically suppress the transcriptional activity of TCR-induced NF-κB and NFAT(*27*), suggesting that sustained TCR signaling is not favored in mature Tregs. This indicates that TCR and IL-2/STAT5 signaling have distinct roles in Treg differentiation and suggests that CD25 acts as a molecular gatekeeper, marking the shift from TCR-dependent to IL-2/STAT5–driven signaling. Biologically, this signal switching may serve to prevent prolonged TCR activation, which could otherwise lead to immune exhaustion or activation-induced cell death.

*Il2ra* transcriptional regulation relies on enhancer-promotor looping(*19*, *20*, *28*). CTCF deletion in mature CD4^+^ and CD8^+^ T cells reduced expression of CD25 following TCR stimulation(*23*). Deleting upstream STAT5-associated super-enhancer elements affects CD25 expression in thymic DN2/DN3 and in mature Tregs(*20*). Several TFs (NF-κB, AP-1, NFAT) along with pioneer factors induced by TCR stimulation (BATF and IRF4) bind predominantly upstream of *Il2ra*(*29*, *30*), suggesting that TCR-associated regulation correlates with upstream enhancer-promoter loop.

Underscoring the potential regulatory effects of intronic elements, three human SNPs associated with CD25 deficiency (rs12722508, rs7909519 and rs61839660) were found within *IL2RA* intron 1(*31*, *32*). These chromatin regions gradually become accessible during Treg development(*20*) and exhibit Treg-specific CpG hypomethylation associated with transcriptional activity that is independent of FOXP3(*33*). In mice, perturbation of cis-regulatory elements with *Il2ra* intron 1 disrupted direct interaction with the TSS and compromised enhancer activity(*18*, *20*). Importantly, both FOXP1 and FOXP3 are capable of binding to the STAT5-associated intronic *IN1m* region(*12*, *34*), and FOXP1 deficiency impairs FOXP3 binding(*12*), suggesting their binding may be cooperative. Collectively, these results emphasize FOXP’s role in sculpting intronic chromatin topology at the *Il2ra* locus.

Although FOXP1 and FOXP3 are the most closely related members of the FOXP family(*35*) and share a majority of their binding sites(*12*), they may individually or cooperatively regulate transcription of various loci, including the *Il2ra* locus. *Il2RA* transcription relies on FOXP3 dimerization, as evidenced by Camperio *et al* who showed that the combination of RelA with wild-type FOXP3 yields maximal *IL2RA* transcriptional activity, as compared with no added activity when RelA was combined with the FOXP3^ΔE251^ 3D-domain swap (3D-DS) mutation(*36*). Similarly, alanine-replacement at M235 within the 3D-DS interface of FOXP3 reduced its DNA binding and FOXP1 interaction, resulting in decreased *Il2ra* expression(*6*). Together, these suggest the involvement of FOXP1 mediated FOXP hetero-dimerization in *Il2ra* regulation. In addition, FOXP3 domain-swapped mutation and Treg ”wannabe” cells did not show a global loss of chromatin contacts(*37*). Structure capture Hi-ChIP results indicated that *Il2ra* is associated with a highly stable and active enhancer that is present in both Treg and Tconv cells, with slightly higher activity in Tregs. Notably, ranked connectivity analysis of FOXP3-specific Hi-ChIP data categorized *Il2ra* as a gene with a “Super Interactive Promoter”(*28*); however, its enhancer-promotor looping intensity is lower than multiple other Treg relevant genes(*28*). These findings support the notion that FOXP1, rather than FOXP3, is the key regulator in establishing the intronic enhancer-promotor looping that is shared in both Treg and Tconv cells. The most likely scenario is that FOXP1 molecules bind DNA through FKHM, after which 3D-DS(*38*, *39*) dimerization bridges DNA together to recruit additional FOXPs (3D-DS or rungs-like(*40*)), in turn stabilizing the nearby DNA and promoting cofactor recruitment.

Our findings provide novel insights into the regulation of *Il2ra* transcription: (1) Temporally, *Il2ra* transcription occurs in two distinct stages: an initiation stage that is independent of FOXP and chromatin topology, and a maintenance stage that depends on the FOXP-associated TAD; (2) Chromosomal looping provides a structural basis for these two transcriptional stages. TCR signals are necessary for the initiation, and IL-2R-mediated signals are required for the maintenance, of high levels of *Il2ra* transcription. (3) A stabilized super-enhancer structure not only confers physical stability and promotes transcriptionally efficient chromatin architecture of the active *Il2ra* locus.

## MATERIALS AND METHODS

### Mice

The generation of *Foxp1* (*Foxp1^fl/fl^*) and *Foxp4* (*Foxp4^fl/fl^*) *loxP*-flanked mice has been previously reported(*9*, *14*, *41*, *42*). *Foxp3* 3’-UTR region (*Foxp3*^YFP-Cre^)(*43*) was obtained from A.Y.Rudensky, *Foxp3^tm9(EGFP/cre/ERT2)Ayr^*/J (*Foxp3*^EGFP-CreERT2^, #016961)*, B6.129P2-Tcrb^tm1Mom^ Tcrd^tm1Mom^/J (Tcrb/Tcrd*^-/-^, #002122), *B6.SJL-Ptprc^a^ Pepc^b^/BoyJ* (CD45.1, #002014) and *B6.PL-Thy1^a^/CyJ* (CD90.1, #000406) transgenic mice were purchased from The Jackson Laboratory. *Rosa26*^lsl-YFP^ mice(*44*) were obtained from F. Costantini (Columbia University) and bred into *Foxp3*^EGFP-CreERT2^ or *Cd4*^CreERT2^ line as a fate-mapper. *Foxp1^fl/fl^* and *Foxp4* (*Foxp1/4^fl/f^*) double-floxed mice were intercrossed with either *Foxp3*^YFP-Cre^, *Foxp3*^EGFP-CreERT2^, *Cd4*^CreERT2^ to generate conditional knockout and tamoxifen-induced conditional knockout in Treg cells. Congenic alleles were bred into *Foxp3*^YFP-Cre^ or *Foxp3*^EGFP-CreERT2^ lines as a distinguishing marker in some competitive experiment settings of this study. All mice used in this study were either purchased on or serially backcrossed for a minimum of 10 generations onto the C57BL/6J background.

All animals were bred and maintained in the Veterans Affairs Palo Alto Health Care System (VAPAHCS) animal facility in an SPF environment at 20°C (68°F) with a 12:12 light-dark cycle. Autoclaved water and an irradiated chow diet (Teklad #2918, Envigo) were provided *ad libitum*. For the longitudinal study in Fig. 1, B and C, 21-day-old control (*Foxp3*^YFP-Cre/X^) and cDKO (*Foxp3*^YFP-Cre/X^ *Foxp1/4*^fl/fl^) female pups were weaned and co-housed until the designated time points. All animal experiment protocols were reviewed and approved by the VAPAHCS institutional animal care and use committee and performed according to the institutional guidelines.

### Chemical administration

To induce Treg-specific knockout in adults, mice underwent oral gavage (P.O.) of 10 µl/g body weight of 2 mg/ml tamoxifen in corn oil each day for 5 consecutive days, and they were analyzed 7 days later, unless noted otherwise. For the BrdU incorporation experiment, mice were given intraperitoneal (I.P.) injections of 10 mg/mL BrdU in 100 uL sterile PBS every 12 hr for 72 hr before sacrifice. BrdU incorporation was measured with the BD BrdU Kit per the manufacturer’s instructions.

### Tissue collection and cell isolation

Tissues from 8-week-old mice, or otherwise noted, were freshly harvested into cold PBS. The thymus, spleen, peripheral lymph nodes (inguinal + brachial + axillary), and Peyer’s patches were dispersed with frosted microscope slides and filtered through 40 µm nylon cell strainers. Erythrocytes were lysed using 10 volumes of RBC lysis buffer. Other non-lymphoid tissues were processed as previously described(*45*). Briefly, tissue samples were chopped and incubated in EDTA-DTT buffer (1 mM EDTA, 1 mM DTT, and 2% FBS in HBSS) at 37 °C for 15 minutes to remove epithelial cells and intraepithelial lymphocytes. After washing, samples were incubated with digestion buffer (30 µg/mL Liberase TM, 50 µg/mL DNase I, and 2% FBS in RPMI-1640) at 37 °C for 45 minutes with gentle shaking every 15 minutes. Pelleted cells after digestion were separated by density gradient and collected at the 40/80% (w/v) Percoll interface.

### Cell culture

Mouse primary T lymphocytes were cultured in complete growth medium (RPMI1640, 10% FBS, 1% L-glutamine, 1% pen/strep, 10 mM HEPES, 1 mM sodium pyruvate, and 0.1 mM 2-mercaptoethanol) at 37°C with 5% CO_2_. CD4^+^ T cells were activated by plate-bound anti-CD3ε mAb (1 µg/mL) and soluble anti-CD28 mAb (1 µg/mL), or anti-CD3/CD28 Dynabeads (Invitrogen).

For iTreg generation, 2 x 10^5^ sorted Tnv cells (CD4^+^CD25^-^YFP^-^CD44^lo^CD62L^hi^) were cultured in each well of a U-bottom 96-well plate (Corning #3799) with 2 ng/mL TGF-β and rmIL2 (100 IU/mL unless otherwise noted).

Ex vivo expansion of murine nTreg cells has been reported previously(*46*). In brief, 1 × 10⁴ sorted nTreg cells (CD4⁺YFP⁺) from Ctrl or cDKO female heterozygous were seeded into individual wells of a 96-well microplate containing CD3/CD28 beads (3:1 ratio) and rmIL-2 (2000 IU/mL) and then restimulated on day 7 with beads at a 1:1 ratio. Cells were harvested at designated time points for analysis by flow cytometry or for use in other downstream experiments.

### Flow cytometry

For general flow cytometry analysis, cells were first blocked with anti-CD16/CD32 and/or 5% rat serum, followed by incubation with Live/Dead viability dye and cell surface markers. For intracellular staining, cells were subsequently fixed and permeabilized with the eBioscience™ FOXP3 staining kit (Invitrogen) according to the manufacturer’s instructions, with minor modifications. Specifically, surface-labeled cells were prefixed with 1% Paraformaldehyde (PFA, w/v) for 10 min, then incubated in Fix/Perm buffer on ice for 20 min, followed by an additional 15 min incubation in permeabilization buffer before adding intracellular antibody cocktails.

To assess pSTAT5 in Tregs while preserving fluorescent protein signals, we adapted a previously described protocol(*47*) with modifications. Briefly, 5x10^6^ lymphocytes were rested in 200 ul RPMI-1640 at 37°C for 30 min, followed by stimulation with IL-2 by combining with 250 ul 2X cytokine stock for an additional 30 min. Immediately after stimulation, samples were fixed by directly adding PFA to a final concentration of 1% and incubated on ice for 10 min. Fixation was then quenched by adding PBS. Pelleted cells were transferred into 96-well microplate and processed as described above. After the 15-minute incubation in permeabilization buffer, cold 90% methanol was added dropwise to the pelleted permeabilized cells, followed by brief mixing and incubation for 10 min. Cells were then rehydrated in 200 μL MACS buffer for 5 minutes, pelleted, and resuspended in intracellular staining antibody cocktails prepared in permeabilization buffer. Intracellular staining was performed for 40 minutes on ice before flow cytometric analysis. Samples were acquired on BD Fortessa or Cytek Aurora flow cytometer. Flow cytometry data were analyzed with FlowJo software (Tree Star Inc.).

### CTV labeling and proliferation assay

2x10^6^ sorted cells or premixed cell mixture were harvested and washed with PBS. The cells were then labeled with 1 uM CellTrace Violet in a 37°C water bath for 30 minutes. The labeling process was stopped by adding 10 times the volume of complete media with 10%FBS. CTV-labeled cells were seeded in anti-CD3ε precoated U-bottom 96-plates (Corning #3799) under iTreg polarizing conditions described above. Cell proliferation was analyzed by flow cytometry and calculated in FlowJo software (Tree Star Inc.)

### Cell viability

Freshly isolated cells were aliquoted into 96-well plates. After staining for surface markers, cell viability was assessed using eBioscience™ Annexin V Apoptosis Detection Kits (Invitrogen) per the manufacturer’s instructions, followed by intracellular staining as described above.

### Cell sorting

Sorting was conducted by using BD AriaIII FACS sorters equipped with a 70-micron nozzle. Cells were sorted into sorting media (Phenol-red free RPMI1640, 2% FBS, 1% L-glutamine, 1% pen/strep, 10 mM HEPES, and 0.1 mM 2-Mercaptoethanol) containing 50% FBS in the FACS or Epi tubes. Post-sort purity was assessed by FACS, confirming that all target cells achieved >98% purity.

To sort out Treg cells containing both GFP and YFP(*Rosa26*) from Tamoxifen-treated mice (icDKO and corresponding Ctrl), a specific filter set was used on SONY MA900 sorter (Sony Biotechnology), GFP and YFP signals were distinguished by applying a 525LP dichroic beam splitter combined with 510/20BP filter for GFP (490-525nm) and a 535/30BP filter for YFP (525-565nm). The sorting was performed with 100 μm microfluidic chips. Bona fide GFP^+^ or YFP^+^ expressed cells were used for compensation. Post-sort purity of GFP^+^YFP^+^ cells achieved > 90%.

### Adoptive T cell transfer

To assess pTreg development under steady-state in vivo, donor naïve T cells (CD4^+^CD25^-^YFP^-^CD44^lo^CD62L^hi^) were double sorted from pooled spleen and PLN to achieve a purity > 99.5%. Tnv from either CD45.2^+^CD90.2^+^ *Foxp3*^YFP-Cre^ *Foxp1/4^+l+^* or *Foxp3*^YFP-Cre^*Foxp1/4^fl/fl^* mice were combined at a 1:1 ratio with congenic CD45.1^+^CD90.2^+^ congenic Tnv cells from *Foxp3*^YFP-Cre^ mice. A total of 1x10^6^ mixed cells in sterile PBS were intravenously infused (I.V.) into sex-matched CD45.2^+^<u>CD90.1</u>^+^ recipients. A portion of pre-transfer cell mixture was fixed and stained on the same day using the same antibody cocktails as for post-transfer analysis. Developed pTreg generated in vivo were harvested 6 weeks after transfer and identified as YFP^+^ cells within donor populations distinguished by unique congenic markers.

To evaluate the in vivo competitiveness of Tregs in the absence of endogenous IL-2, a total of 2×10^5^ cells, derived from ex vivo expanded CD4^+^YFP^+^ nTregs from CD45.2^+^CD90.2^+^ *Foxp3*^YFP-Cre^ or *Foxp3*^YFP-Cre^*Foxp1/4*^fl/fl^, were mixed with CD45.2^+^CD90.1^+^ *Foxp3*^EGFP-CreERT2^ at a 1:1 ratio and intravenously injected into CD45.1^+^ Tcr*b/Tcrd*^-/-^ recipients (fig. S7). Samples were harvested 14 days after the adoptive transfer, and donor-derived cell populations were distinguished by congenic markers. The experimental endpoint was determined by the assigned time point and health status of the animals, per protocols approved by the IACUC.

### Bulk RNA sequencing and annotation

FOXP3^+^ committed Treg cells (CD4^+^YFP^+^) were sorted from pooled spleen and PLN of 8-week-old *Foxp3*^YFP-Cre/X^*Foxp1/4^+/+^* or *Foxp3*^YFP-Cre/X^*Foxp1/4^fl/fl^* heterozygous. RNA isolation and sequencing were performed as previously reported(*15*). Sequencing data were aligned and annotated to the mouse genome assembly GRCm38.p6/mm10 at Stanford Genomic Core.

Individual raw fastq files underwent quality assessment using FastQC (v 0.12.1). Adapter and low-quality sequences at the end of each read were removed from paired-end reads using Trimmomatic v0.39 with TRAILING=30. Trimmed reads were further mapped to the mouse reference genome (GRCm38.p6/mm10) using Burrow-Wheeler Aligner (BWA-MEM, v0.7.17). The raw counts and gene-level counts were calculated using featureCounts in Subread package v2.0.6. A prefilter was done to the dataset to remove genes with < 10 counts per row. Differential gene expression analysis for each comparison was conducted on the raw counts data using DESeq2 v1.40.2 in R v4.3.3. A significantly differentially expressed genes were considered under FDR < 0.05.

### Bulk ATAC sequencing and peak calling

Lymphocytes were collected and pooled from spleen and PLN from 8-week-old mice. Cells were enriched by magnetically negative selection for CD4^+^ cells, then sorted out CD4^+^YFP^+^ cells on BD AriaIII on a 70-micron nozzle, into 200 ul sorting medium containing 50% FBS in Epi tubes. Approx. 30,000-50,000 Treg cells can be obtained from each mouse.

ATAC-seq preparation/transposition reaction was performed as previously described(*48*) with minor modifications. In brief, cells were collected and then lysed in 100 μL cold ATAC-RSB buffer (0.5% NP40, 0.1% Tween20, and 0.1% Digitonin). Pellet nuclei were collected at 550rcf for 30min at 4 °C in a fixed angle centrifuge, followed by adding 50 μL transposition mixture (Nextera Kit, Illumina Cat # FC-121-1030) and incubating at 37 °C for 30 min in a thermomixer (Eppendorf, Thermomixer F1.5) with 1000 rpm mixing. Post-transposition DNA fragments were purified by using the Qiagen MinElute PCR Purification Kit and protocol (Cat# 28004).

For library preparation, samples were amplified by QPCR (NEB Next High-Fidelity 2X PCR master mix) with Nextrra Primers with adaptors (table S4). Post-amplification cleanup was performed by double-sized SPRI beads selection (Beckman Coulter, B23317). Amplified Library quality was confirmed on Agilent Tapestation D1000 and analyzed on Tapestation 4200 (Agilent). Sequencing was performed on HiSeq 4000, 2x75 pair-ends at Stanford Genomics Service Center (SGSC).

ATAC-seq data was aligned to the mouse reference genome (GRCm38.p6/mm10) using Bowtie2 v2.5.1 after the same quality assessment as the RNAseq data. Mitochondrial reads, duplicates, low-quality alignments, multi-mapped reads, ENCODE blacklist regions were removed using SAMtools v1.18 and bedtools v2.31.0. A shift read coordinates was performed using alignmentSieve function in deepTools v3.3.0 to correct for the potential small DNA insertion bias. Peaks were called using MACS2 v2.2.7.1 in its narrow peak mode for paired-end reads. The non-reproducible peaks in regions of low coverage and other artifacts were removed across the biological replicates using bedtools v2.31.0, and consensus peaks were further merged peak sets using BEDOPS v2.4.41. Peaks with Fraction of Reads in Peaks (FRiP) less than 0.2 were removed, and remaining reads were summarized within each peak using featureCounts in the Subread package v2.0.6. The retained peaks were further annotated using TSS annotations from Ensembl rather than from the UCSC-based (TxDb.Mmusculus.UCSC.mm10.knownGene) and *org.M. Mm. eg, b* in R.

### Bioinformatics data analysis

Differentially expressed genes were identified using DESeq2 as previously described. Genes showing significant differential expression (FDR < 0.05) and a log2-fold change greater than 0.5 or less than -0.5 were analyzed separately for GSEA against the different MSigDB Gene Sets from *msigdbr* v7.5.1 using *clusterProfiler* v4.10.0 and only gene sets meeting a Benjamini-Hochberg adjusted FDR threshold of less than 0.05 were retained. The GO enrichment analysis was performed for all subontologies against the Genome wide annotation for Mouse from *org.Mm.eg.db* v3.18 using *clusterProfiler* v4.10.0 and only GO terms meeting a q-value threshold of less than 0.05 were retained. Pathway enrichment analysis was restricted to KEGG and only pathways meeting a Benjamini-Hochberg adjusted FDR threshold of less than 0.1 were retained.

Publicly accessible sequencing data and Hi-C matrix files used in this study are listed in table S3. Raw Sequence Read Archive (SRA) files were filtered, trimmed using sra-tools v3.1, and realigned to the mouse reference genome (GRCm38.p6/mm10) using Bowtie2. Peak or region visualization were performed using Integrated Genome Viewer (IGV), with grouped scaling and normalization based on total mapped reads. For in situ Hi-C, followed by assigned alignments to MobI fragments. Contact matrices were generated through HiC-pro pipeline (v3.1.0)(*49*) on SLURM cluster. Valid interaction products were normalized using the ICE algorithm(*50*). Contact maps were visualized using HiGlass (https://github.com/higlass/higlass-docker) running in Docker v4.36.0.

For in silico Hi-C, the model was trained using Hi-C matrix, CTCF ChIP-seq, and ATAC-seq from wild-type mouse Tregs, preprocessed according to Seq-N-Slide pipeline. A multimodal neural network architecture integrating DNA sequence and epigenomic features through parallel convolutional and transformer layers was optimized using the Adam optimizer (learning rate 1×10^-4^) with a cosine learning rate scheduler (100-epoch period). Training employed a batch size of 8 and was conducted on NVIDIA Tesla GPU clusters. The minimal validation loss was achieved at 24 epochs. For de novo prediction, 2-Mb genomic regions were analyzed using reference DNA sequence(mm10), CTCF ChIP-seq, and ATAC-seq from Tregs as input. Insulation scores and Virtual 4C profiles were calculated as previously described(*51*).

### Chromatin immunoprecipitation

YFP^+^ Tregs were sorted and harvested from 8-week-old *Foxp3*^YFP-Cre^*Foxp4^+l+^*or *Foxp3*^YFP-Cre^*Foxp4^fl/fl^* mice. The cells were fixed with 1% paraformaldehyde for 10 minutes at room temperature to crosslink proteins and associated DNA, followed by quenching with 125 mM ice-cold glycine. After double-washing, the cell pellets were resuspended in 200 µL Pierce RIPA buffer (Thermo Scientific #89900) containing protease inhibitors (Roche) and incubated at 4°C overnight. The lysate pellets were sonicated at 4°C in 1.5 mL microtubes using Bioruptor Pico (Hologic Diagenode) with a 15-second on/90-second off cycle for a total of 8 cycles. The resulting DNA fragments were approximately 200-500 bp, as confirmed by agarose gel electrophoresis. To each sample, 1 µg of BS-3 crosslinker (Abcam) conjugated H3K27ac-Dynabeads Protein G complex was added and incubated at 4°C overnight on a shaker. Following incubation, the beads were enriched using a magnetic rack and washed twice with ice-cold RIPA buffer. Reverse crosslinking was performed by incubating the samples in RIPA buffer with proteinase K (200 µg/mL) at 65°C for 2 hours while shaking at 500 rpm in a thermomixer. The released DNA fragments were collected and purified using the QIAGEN QIAquick PCR Purification Kit and then reconstituted in TE buffer.

Primers were designed using Primer3 and are as follows: *mIN1m* Forward: 5’-GTTGTGGTGGCTTTCCCTGC; *mIN1m* Reverse: 5’-GTGTCACTAGCACGGGGACA; *mIN1b* Forward: 5’-CAGAAGACTGTGGGCGGGAT; *mIN1b* Reverse: 5’- GATTTCTTGGGAGGGGGCCA. Primers for *Gapdh* were included in the ChIP-IT qPCR analysis kit (Active Motif, #53029), and copy numbers were calculated according to the manufacturer’s instructions.

### Quantitative real-time PCR

Total RNA was extracted using the RNeasy Micro Kit (Qiagen) according to the manufacturer’s instructions. cDNA was reverse transcribed with the Superscript III first strand synthesis system (Invitrogen) and amplified with TaqMan Universal Master Mix (Applied Biosystem). Gene expression was quantified using TaqMan probes (KEY RESOURCE TABLE). Ct values were normalized to the mean Ct of housekeeping gene *Actb*, and relative fold changes were calculated using the 2-ΔΔCt method.

### Quantification and statistical analysis

FlowJo v10.10.0 (BD), Prism v10.4.1(GraphPad), R v4.3.3, and RStudio were used for statistical analysis. Normal distribution was assumed for samples, a two-tailed non-parametric Mann-Whitney U test was used to compare between two groups, and an ordinary one-way ANOVA with adjusted Tukey’s post-hoc test was performed for multiple comparisons. *P*-values < 0.05 were considered statistically significant. All graphs show averages of the mean ± standard error of mean unless otherwise stated.

## Supporting information

Supplemental Tble 2

## Supplementary Materials

Fig. S1 to S10

Table S1 to S5

## Other Supplementary Material for this manuscript includes the following

Data file S1 (will provide upon acceptation) MDAR Reproducibility Checklist

## Acknowledgments

We thank David Hildeman and Fadi Lakkis for their critical reading and helpful suggestions and Hong Zhen and Zachary Joseph for assistance with bioinformatic analyses.

## Author contributions

Conceptualization and methodology: D.D., L.E.H., and J.S.M.

Investigation: D.D., J.Z., J.P. and Y.K.

Writing – original manuscript: D.D.

Writing – review & editing: D.D., L.E.H., J.X.L., W.J.L. and J.S.M.

Supervision: J.S.M.

Funding acquisition: J.S.M.

## Data and materials availability

- RNA-seq data (GSE300286) and ATAC-seq data (GSE307753) have been deposited at the Gene Expression Omnibus (GEO) database and are publicly available as of the date of publication.
- This study analyzes publicly accessible data. The accession numbers for those datasets are listed in the table S3.
- Custom code used in this study has been deposited at Zenodo: https://doi.org/10.5281/zenodo.17823014.
- Any additional information required to reanalyze the data reported in this paper is available from the lead contact upon request.

---

**fig. S1.**
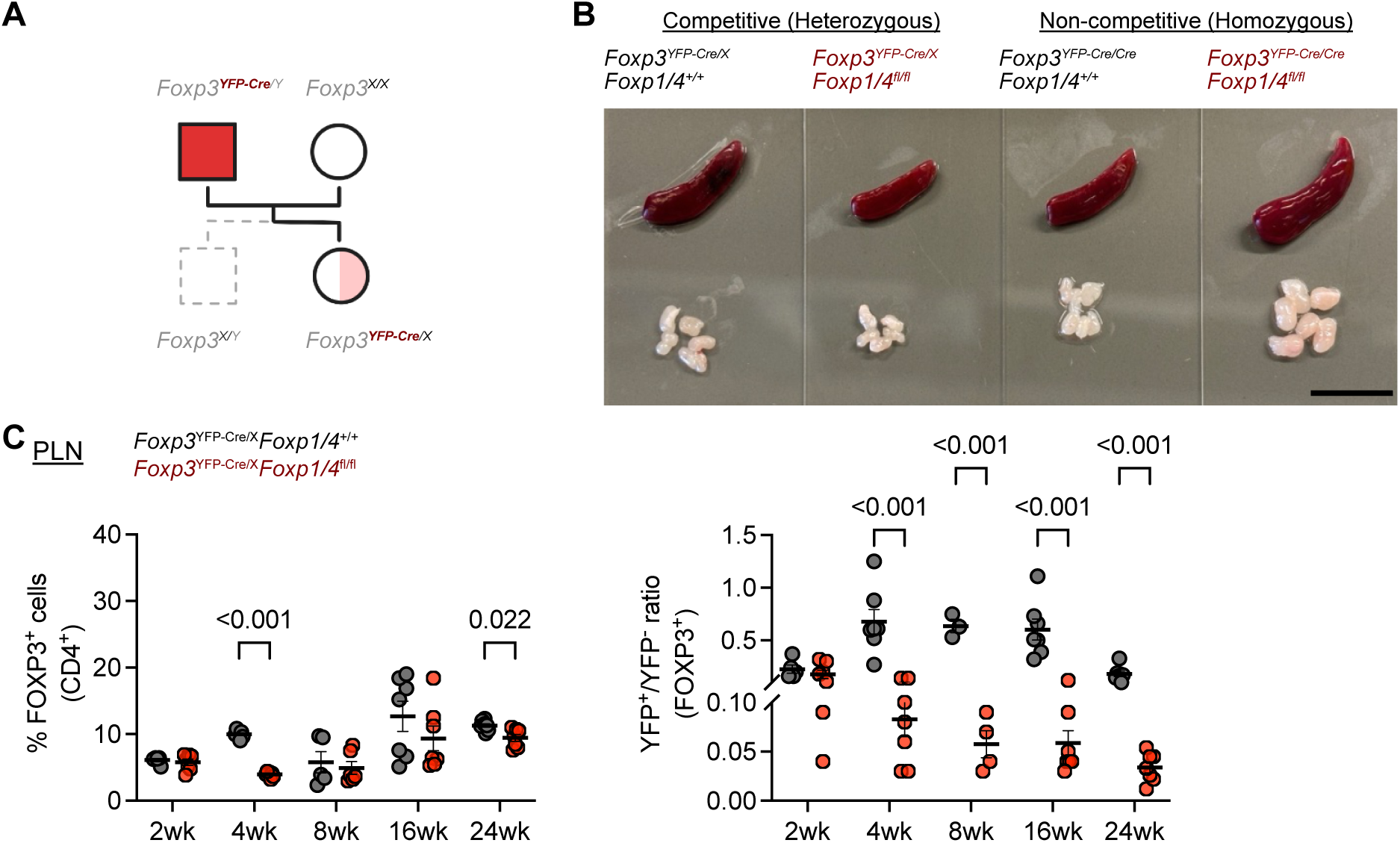
Breeding and phenotypic assessment of *Foxp3^YFP-Cre/X^Foxp1/4^fl/fl^* mice. (A) Breeding strategy of generating *Foxp3^YFP-Cre/X^Foxp1/4^fl/fl^* female heterozygotes. (B) A representative appearance of spleen and peripheral lymph nodes from *Foxp3^YFP-Cre/X^Foxp1/4^fl/fl^* heterozygotes and *Foxp3^YFP-Cre/Cre^Foxp1/4^fl/fl^* homozygous at 8 weeks of age. (Scale bar = 1cm). (C) The frequency of FOXP3^+^ Treg within CD4^+^ cell and the YFP^+^/YFP^−^ ratio at the indicated time points (n= 5 to 8 per group). Data are presented as mean ± SEM, each dot represents data from an individual mouse. Statistical analysis: two-tailed unpaired t-test, *P* values are shown for significant difference (*P* < 0.05).

**fig. S2.**
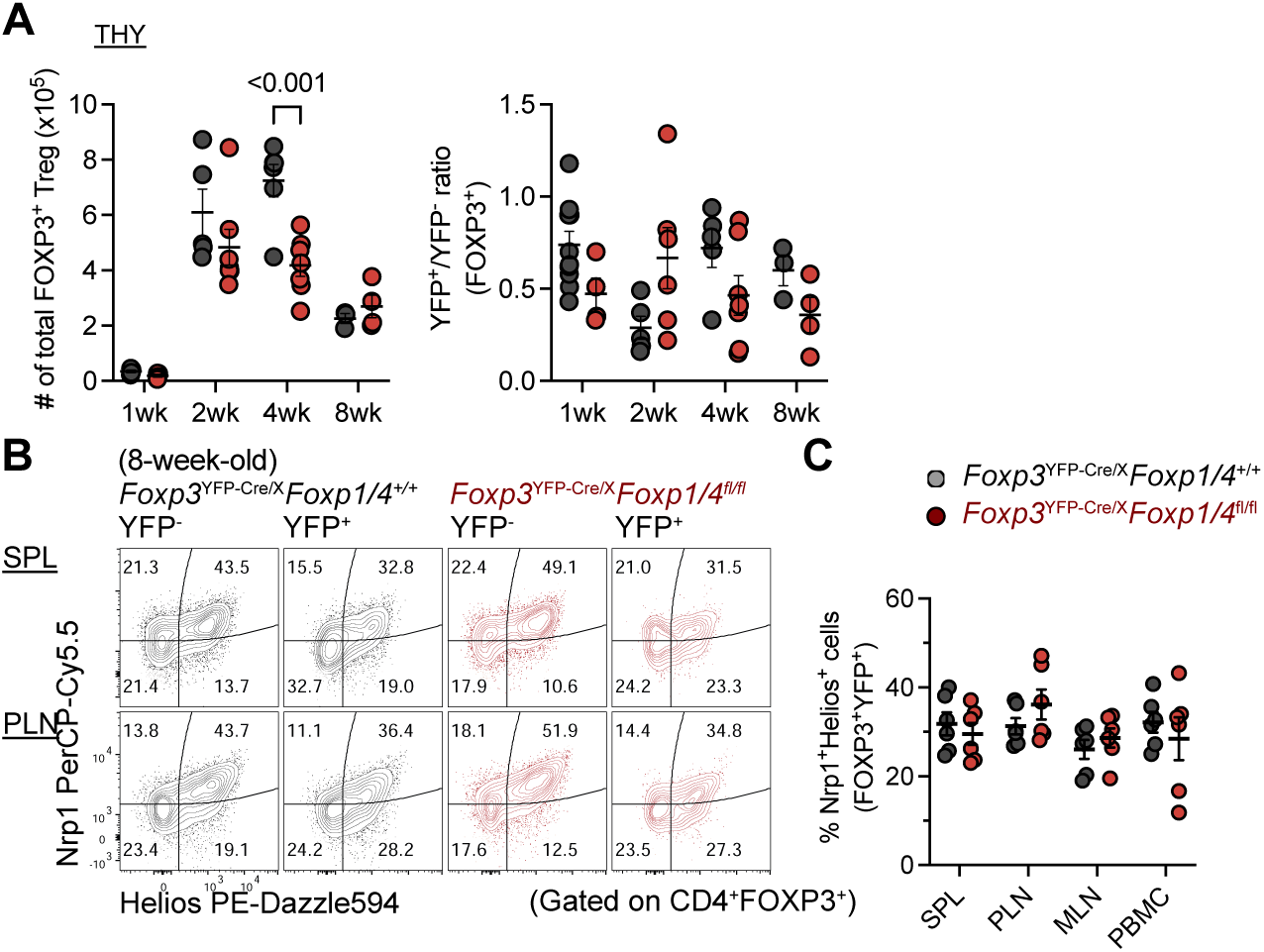
Loss of FOXP1/FOXP4 does not affect tTreg development or peripheral seeding. (A) Total number and YFP^+^/YFP^-^ ratio of Tregs in thymus of mice at 1, 2, 4 and 8 weeks of age (n = 4-10 per group). (B and C) Representative flow cytometry plots and quantitation illustrate the proportion of the tTreg (Nrp1^+^Helios^+^) compartment in the splenic lymphoid organs (SLOs) of 8-week-old mice. Statistical analysis: two-tailed unpaired t-test, *P* values are shown for significant difference (*P* < 0.05).

**fig. S3.**
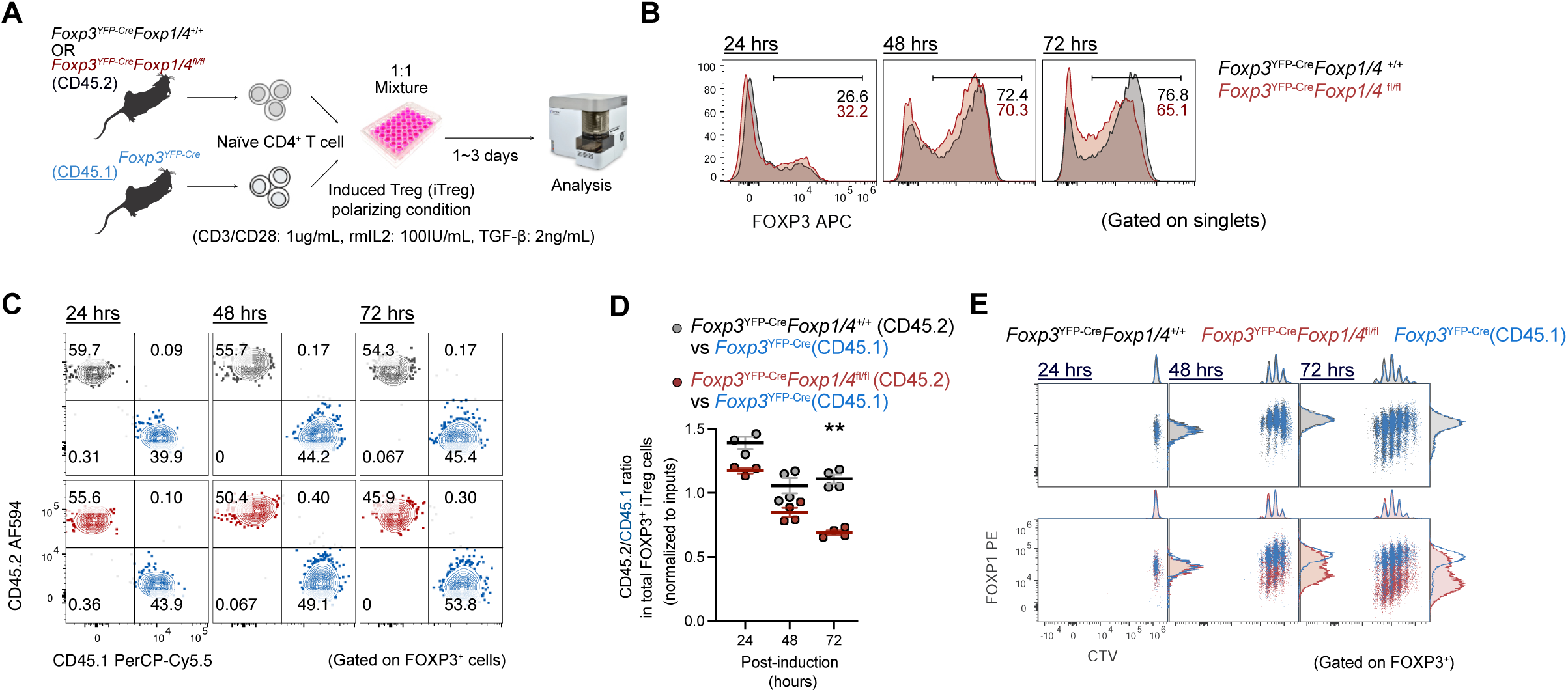
FOXP1/FOXP4-deficient iTreg develop and proliferate normally. (A) Schematic workflow for studying *in vitro* iTreg polarization under competitive condition. (B) Proportion of newly generated FOXP3^+^ iTreg cells (boxed quadrant) during the first 24 and 48 hours of polarization. (C) Representative flow cytometric plots of iTreg competitiveness after induction, and (D) quantification of their relative ratio within FOXP3^+^ cells (n=3 to 4 per group, representative of two independent experiments). (E) CTV-labeled tracking of cell proliferation was measured at 24-hour intervals during competitive co-culture iTreg polarization. Staked histogram showing FOXP1 expression level and cell proliferation during iTreg induction. Statistical analysis: two-tailed unpaired t-test, \*\**= P* <0.01.

**fig. S4.**
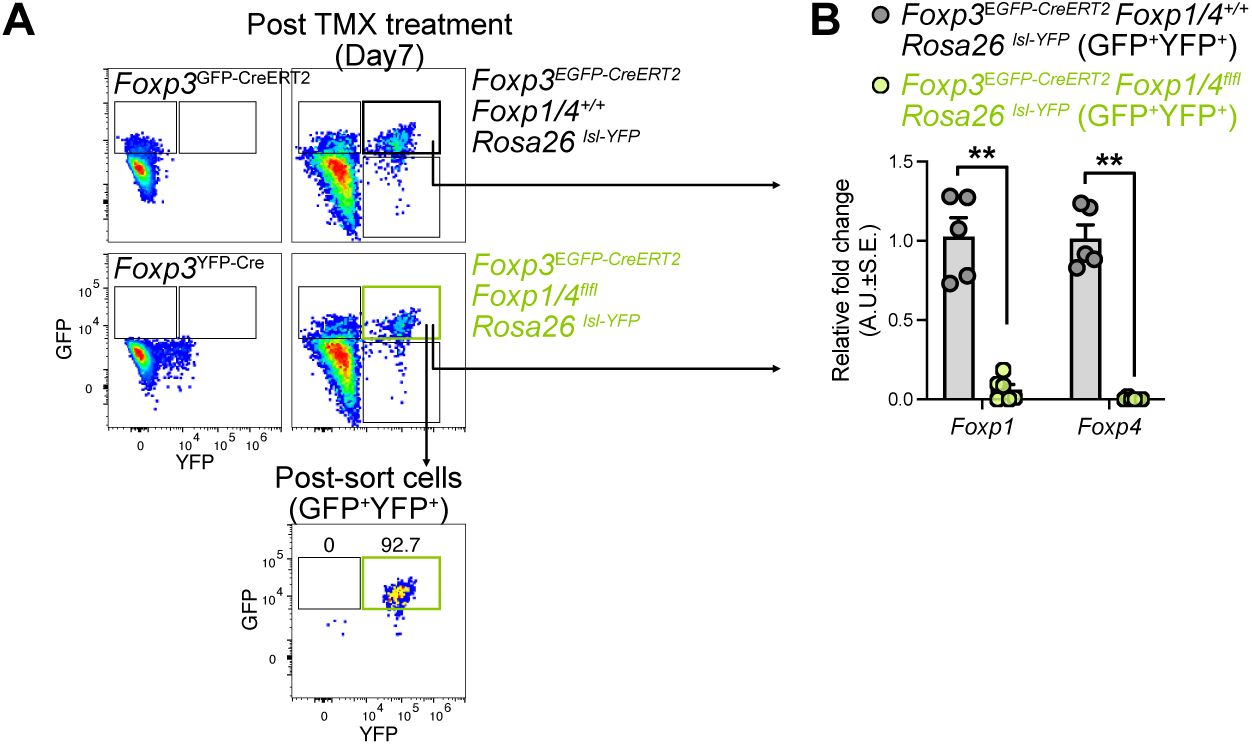
Acute depletion of FOXP1 and FOXP4 in Tregs. (A) Representative flow cytometric plots of sorted YFP^+^ cells from GFP^+^ Tregs. (B) Quantification of relative *Foxp1* and *Foxp4* transcripts in sorted YFP^+^GFP^+^ cells seven days post-inducible knockout. Data are presented as mean ± SEM, each dot represents data from an individual mouse. Statistical analysis: two-tailed unpaired t-test, ** *= P* <0.01

**fig. S5.**
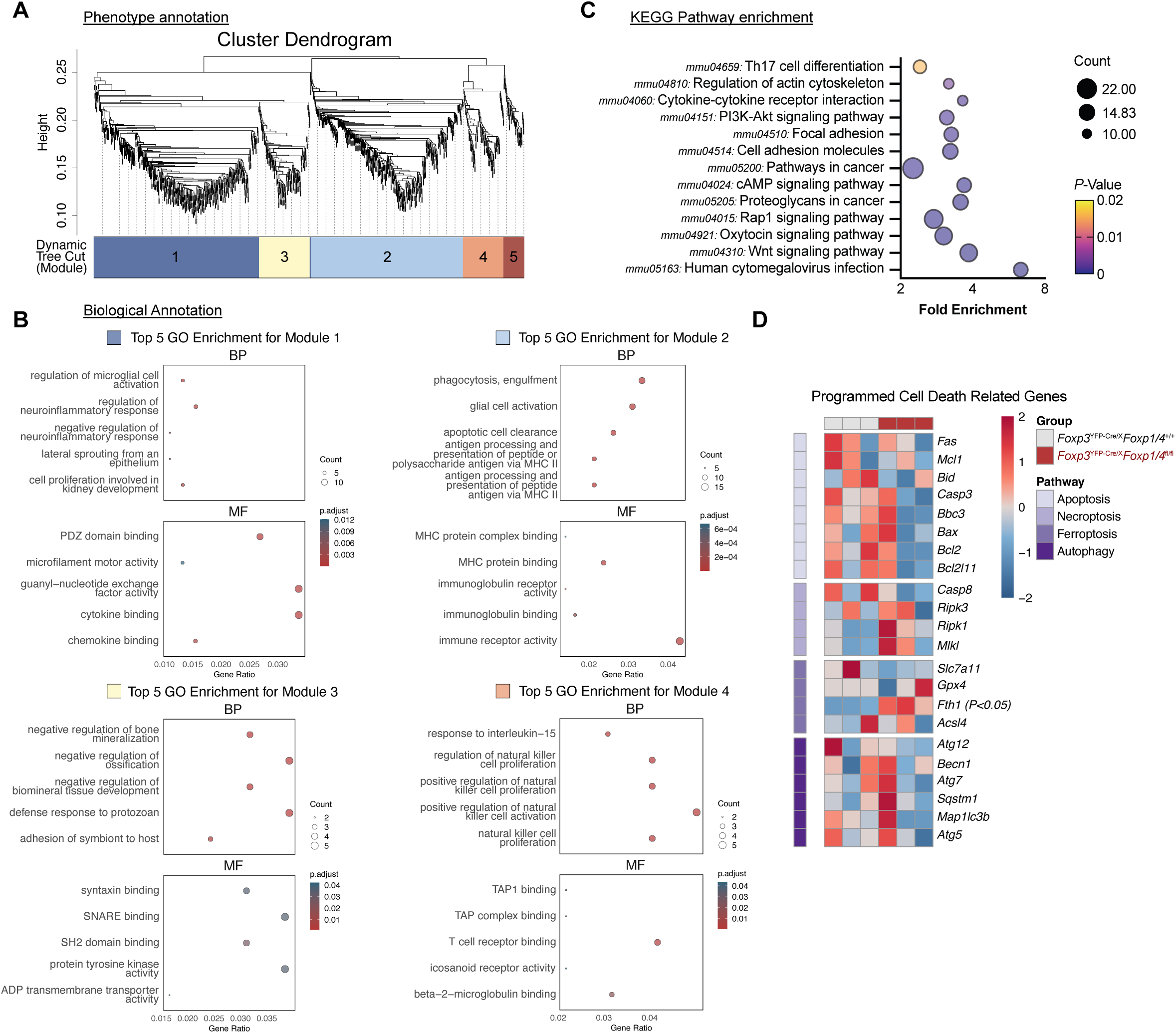
Transcriptomic annotation of FOXP1/FOXP4 deficient Tregs. (A) Dendrogram of DEGs from RNA-seq data clustered by weighted gene co-expression network analysis, using hierarchical clustering (McQuitty method) and dynamic tree cutting based on TOM dissimilarity. (B) Biological function annotation (Biological Process; BP and Molecular Function; MF) of the top 4 clustered modules identified in (A). (C) KEGG pathway enrichment analysis of genes shared between RNA-seq and ATAC-seq datasets. (D) Heatmap of transcription level of programmed cell death-related genes.

**fig. S6.**
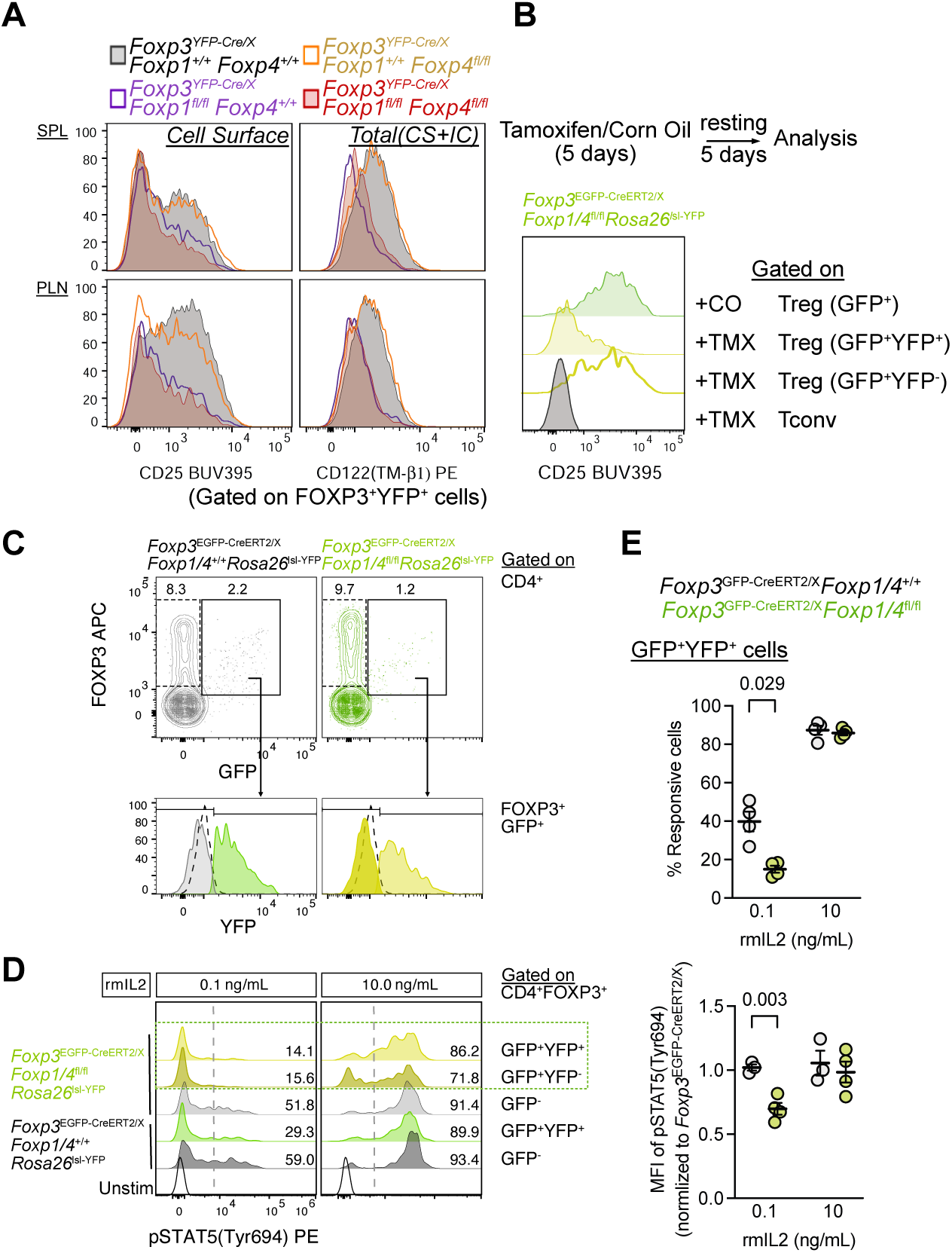
FOXP1 is required for competition for IL-2 responsiveness and is associated with CD25. (A) Histogram of cell surface CD25 and total CD122 (cell surface plus intracellular; CS+IC) expression level in FOXP3⁺YFP⁺ cells from all four strains. (B) Stacked histogram of cell surface CD25 in *Foxp3-*driven FOXP1/FOXP4 acutely depleted Treg cells. (C and D) Representative flow cytometric plots of gating strategy (C) and pSTAT5_Tyr694_ levels in PLN Tregs obtained from FOXP1/FOXP4 acutely depleted mice (D). (E) Quantification of the proportion of IL-2-responsive cells and relative pSTAT5_Tyr694_ levels in Tregs from (D) (n = 4 per group). Data are presented as mean ± SEM, each dot represents data from an individual mouse. Statistical analysis: two-tailed unpaired t-test, exact *P* values are shown for significant difference (*P* < 0.05).

**fig. S7.**
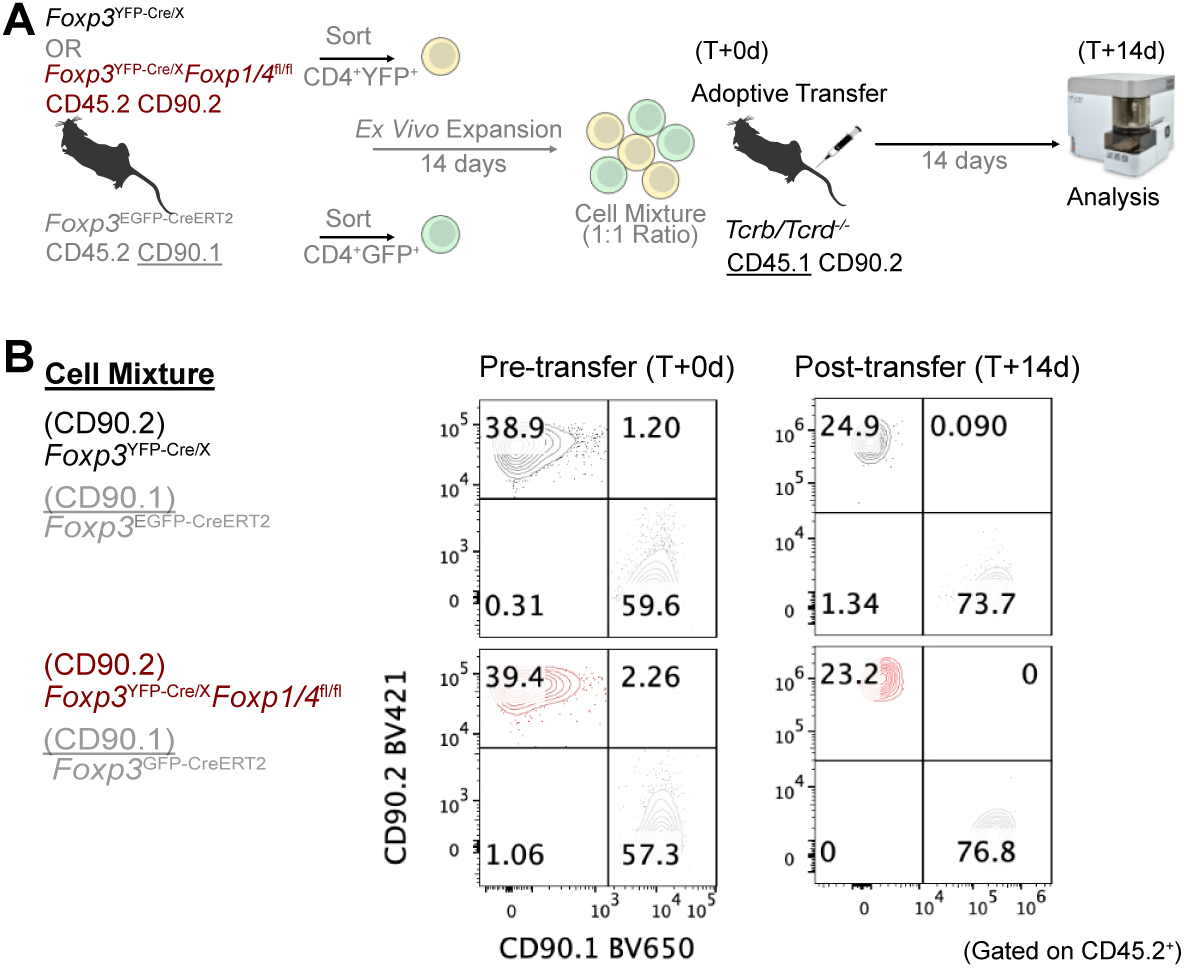
No competitive disadvantage of FOXP1/FOXP4-deficient Tregs in the absence of conventional T cells. (A) Schematic of workflow to assess Treg competitiveness for IL-2 *in vivo* by using *ex vivo* expanded Tregs. (B) Representative flow cytometric plots showing the relative competitiveness of adoptively transferred donor cells in congenic *Tcrb/Tcrd^-/-^* hosts.

**fig. S8.**
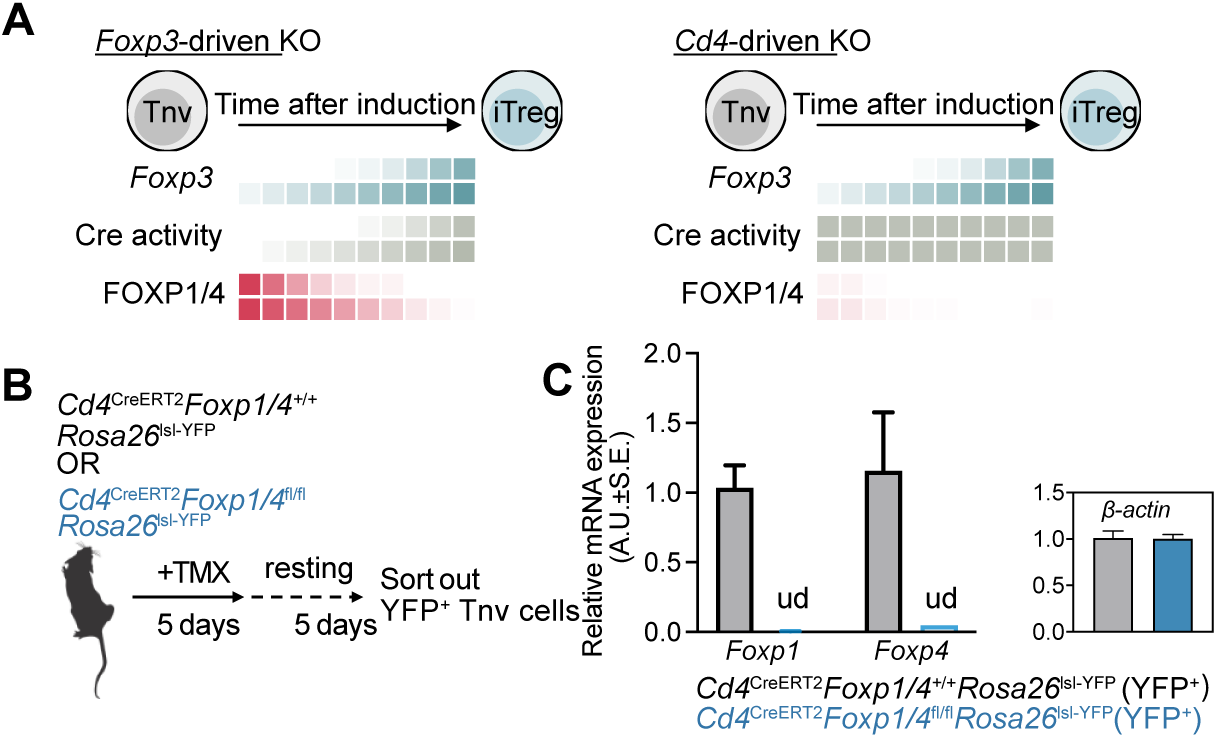
CD4^Cre-ERT2^-mediated deletion of *Foxp1/4*. (A) Cartoon illustration showing how FOXP1/FOXP4 levels change during iTreg polarization under the control of *Foxp3*-driven or *Cd4*-driven Cre activity. (B) Workflow of obtaining *Foxp1/4*-deficient Tnv from *Cd4*^CreERT2^ mice. (C) Quantification of *Foxp1* and *Foxp4* knockout efficiency in *Cd4*-driven inducible knockout models.

**fig. S9.**
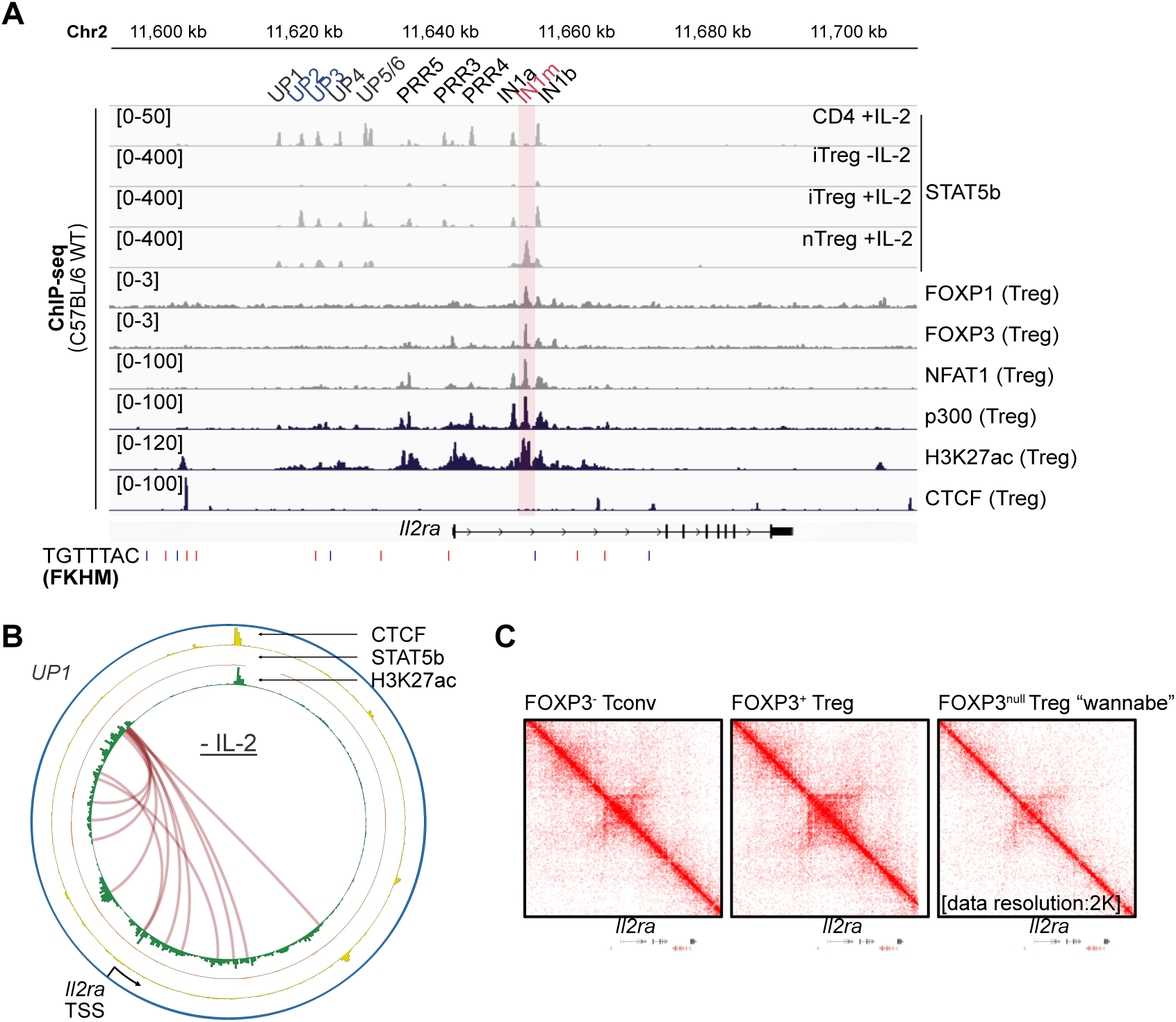
Regulatory landscape and chromatin architecture of Il2ra locus in Treg. (A) STAT5b ChIP-seq tracks at the *Il2ra* locus during Treg development, and ChIP-seq of FOXP1, FOXP3, NFAT1, P300, H3K27ac, and CTCF at the *Il2ra* locus in wildtype mature Tregs. Reanalyzed from publicly accessible datasets (Supplementary Table 3). Red shading represents the IN1m region of the *Il2ra* locus. (B) RNA-PolII ChIA-PET plot of chromatin interactions in the *Il2ra* locus of wild-type CD4 cells, without IL-2 stimulation. (C) In-situ Hi-C plots at the *Il2ra* locus in Tconv, Treg from spleen and Treg “wannabe” (samples GSM6705669, GSM6705671, and GSM6705673).

**fig. S10.**
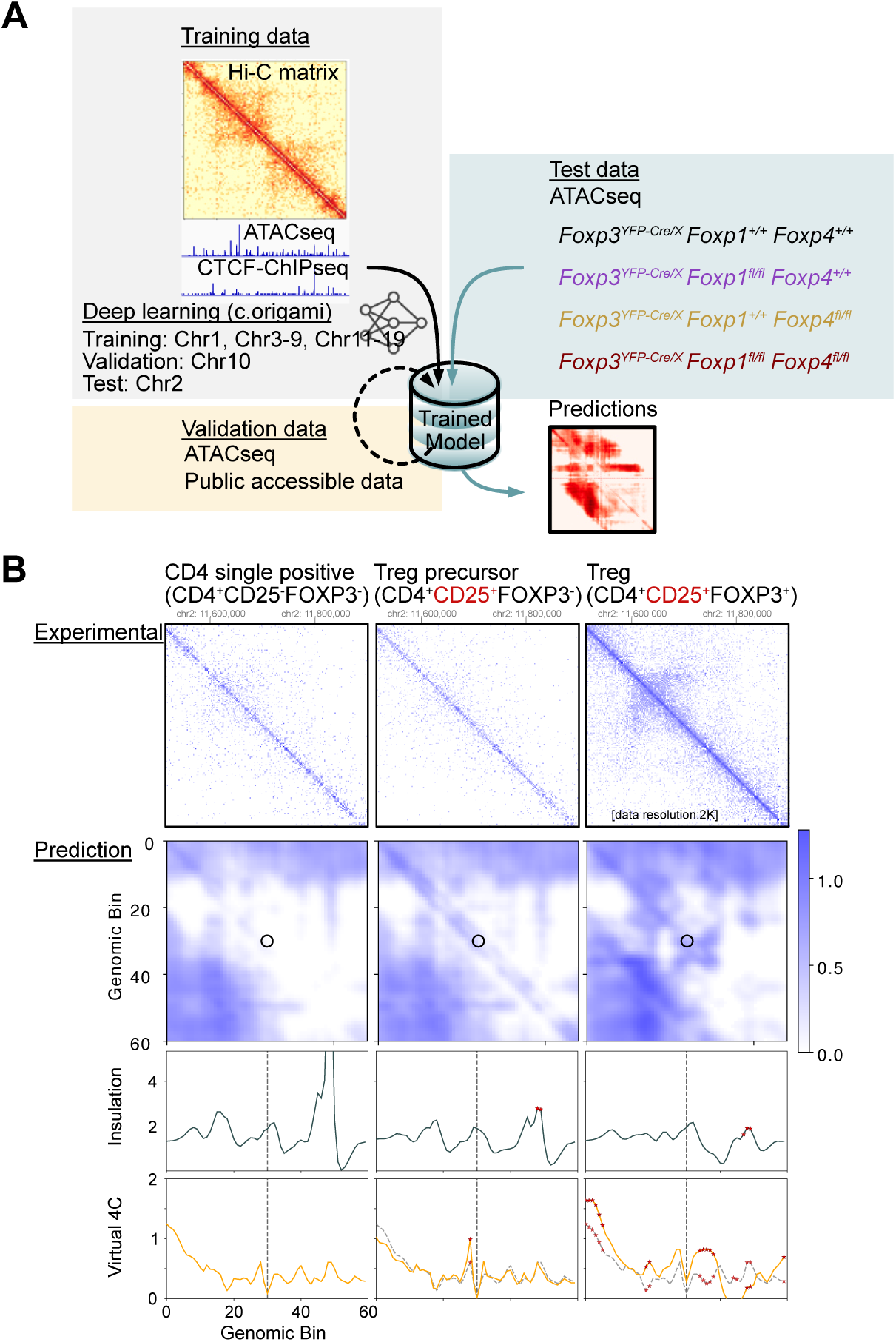
Treg-specific trained model for *in silico* Hi-C analysis. (A) Schematic workflow for *in silico* Hi-C prediction in mouse Tregs using publicly accessible data (Hi-C:GSM6705675, CFCF-ChIP: GSM7213946, ATAC-seq: GSM9230397). (B) Validation of the predictive performance of the trained model. Experimental data were obtained from in situ Hi-C of thymic Tregs, the prediction matrix was generated based on analysis of ATAC-seq from thymic Tregs (SRR5385309, SRR5385308, and SRR5385307). Insulation score and virtual 4C profiles calculated to predict TAD formation at *Il2ra* locus. Open circle on the matrix and the vertical line on both the insulation score and virtual 4C indicates the *Il2ra* TSS.

