## Supplemental Tables 3-5 for "FOXP-stabilization of the *Il2ra* super-enhancer structure augments Treg fitness"

**Table S3. List of publicly available data used in this study, related to Fig. 6, fig. S9 and S10.**

| File Identity | BioSample | SeqType | Database |
|---|---|---|---|
| SRR5385310 | DP | ATAC | DDBJ |
| SRR5385309 | ImCD4SP | ATAC | DDBJ |
| SRR5385308 | CD25+tTreg prec | ATAC | DDBJ |
| SRR5385307 | tTreg | ATAC | DDBJ |
| SRR5385355 | DP, H3K27ac ChIP-Seq | ChIP-seq | DDBJ |
| SRR5385353 | ImCD4SP, H3K27ac ChIP-Seq | ChIP-seq | DDBJ |
| SRR5385351 | CD25+tTreg prec, H3K27ac ChIP-Seq | ChIP-seq | DDBJ |
| SRR5385349 | tTreg, H3K27ac ChIP-Seq | ChIP-seq | DDBJ |
| GSM7165880 | WT Treg-1, H3K27ac ChIP-Seq | ChIP-seq | GEO |
| GSM7165882 | WT Treg-2, H3K27ac ChIP-Seq | ChIP-seq | GEO |
| GSM7165890 | WT Treg, NFAT1 ChIP-Seq | ChIP-seq | GEO |
| GSM7165888 | WT Treg, p300 ChIP-Seq | ChIP-seq | GEO |
| GSM7213946 | Treg, CTCF ChIP-seq | ChIP-seq | GEO |
| GSM3430729 | Foxp1+ Treg, rep1, FOXP1 ChIP-seq | ChIP-seq | GEO |
| GSM3430735 | Foxp1+ Treg, rep1, FOXP3 ChIP-seq | ChIP-seq | GEO |
| GSM3430737 | Foxp1- Treg, rep1, FOXP3 ChIP-seq | ChIP-seq | GEO |
| GSM2734699 | IN1 CD4T +IL2, H3K27Ac ChIP-seq | ChIP-seq | GEO |
| GSM2734700 | IN2 CD4T +IL2, H3K27Ac ChIP-seq | ChIP-seq | GEO |
| GSM8661318 | iTreg, Cont, STAT5B ChIP-seq | ChIP-seq | GEO |
| GSM8661319 | iTreg, IL2, STAT5B ChIP-seq | ChIP-seq | GEO |
| GSM8661334 | ex vivo Treg, STAT5B ChIP-seq | ChIP-seq | GEO |
| GSM2734684 | CD4T, +IL2, STAT5B ChIP-seq | ChIP-seq | GEO |
| GSM6705657 | CD4-positive, alpha-beta thymocyte [Thy] | in situ HiC | GEO |
| GSM6705655 | Treg precursor - CD25+ Foxp3- [Thy] | in situ HiC | GEO |
| GSM6705675 | Treg - thymus - Fox3p-GFP+ [Thy] | in situ HiC | GEO |
| GSM6705673 | "wannabe" Treg - thymus - Foxp3-KO [Thy] | in situ HiC | GEO |
| GSM6705669 | Tcon-SPL | in situ HiC | GEO |
| GSM6705671 | Treg-SPL | in situ HiC | GEO |

**Table S4. HiSeq pools and adapters for ATAC-seq library preparation, related to Fig. 4.**

| *Pool* | *Samples* | *Index* | *Sequence* |
|---|---|---|---|
| **A** | 9084 | Ad2.4_TCCTGAGC | CAAGCAGAAGACGGCATACGAGATGCTCAGGAGTCTCGTGGGCTCGGAGATGT |
| | 9085 | Ad2.6_TAGGCATG | CAAGCAGAAGACGGCATACGAGATCATGCCTAGTCTCGTGGGCTCGGAGATGT |
| | 9086 | Ad2.7_CTCTCTAC | CAAGCAGAAGACGGCATACGAGATGTAGAGAGGTCTCGTGGGCTCGGAGATGT |
| | 9106 | Ad2.1_TAAGGCGA | CAAGCAGAAGACGGCATACGAGATTCGCCTTAGTCTCGTGGGCTCGGAGATGT |
| | 9186 | Ad2.9_GCTACGCT | CAAGCAGAAGACGGCATACGAGATAGCGTAGCGTCTCGTGGGCTCGGAGATGT |
| | 9196 | Ad2.11_AAGAGGCA | CAAGCAGAAGACGGCATACGAGATTGCCTCTTGTCTCGTGGGCTCGGAGATGT |
| | 9197 | Ad2.12_GTAGAGGA | CAAGCAGAAGACGGCATACGAGATTCCTCTACGTCTCGTGGGCTCGGAGATGT |
| **B** | 9084 | Ad2.5_GGACTCCT | CAAGCAGAAGACGGCATACGAGATAGGAGTCCGTCTCGTGGGCTCGGAGATGT |
| | 9107 | Ad2.2_CGTACTAG | CAAGCAGAAGACGGCATACGAGATCTAGTACGGTCTCGTGGGCTCGGAGATGT |
| | 9108 | Ad2.3_AGGCAGAA | CAAGCAGAAGACGGCATACGAGATTTCTGCCTGTCTCGTGGGCTCGGAGATGT |
| | 9188 | Ad2.10_CGAGGCTG | CAAGCAGAAGACGGCATACGAGATCAGCCTCGGTCTCGTGGGCTCGGAGATGT |
| | 9199 | Ad2.13_GTCGTGAT | CAAGCAGAAGACGGCATACGAGATATCACGACGTCTCGTGGGCTCGGAGATGT |
| | 9176 | Ad2.14_ACCACTGT | CAAGCAGAAGACGGCATACGAGATACAGTGGTGTCTCGTGGGCTCGGAGATGT |

**Table S5. Key resource.**

| REAGENT or RESOURCE | SOURCE | IDENTIFIER |
|---|---|---|
| **Antibody** | | |
| BD Pharmingen™ APC Rat Anti-Mouse CD62L (MEL-14) | BD biosciences | 553152 |
| BD Pharmingen™ Alexa Fluor® 647 Mouse anti-BrdU (3D4) | BD biosciences | 560209 |
| BD Horizon™ BUV395 Rat Anti-Mouse CD25 (PC61) | BD biosciences | 564022 |
| BD Horizon™ BUV496 Hamster Anti-Mouse CD3ε (145-2C11) | BD Biosciences | 612955 |
| Brilliant Violet 785™ anti-mouse CD3 (17A2) | Biolegend | 100232 |
| PE anti-mouse CD4 (GK1.5) | Biolegend | 100408 |
| Alexa Fluor® 700 anti-mouse CD4 (GK1.5) | Biolegend | 100430 |
| Pacific Blue™ anti-mouse CD5 (53-7.3) | Biolegend | 100641 |
| PerCP/Cyanine5.5 anti-mouse CD8a (53-6.7) | Biolegend | 100734 |
| PE/Cyanine7 anti-mouse CD25 (3C7) | Biolegend | 101916 |
| PE anti-mouse CD25 (PC61) | Biolegend | 102008 |
| Ultra-LEAF™ Purified anti-mouse CD28 (37.51) | Biolegend | 102116 |
| Pacific Blue™ anti-mouse/human CD44 (IM7) | Biolegend | 103020 |
| Pacific Blue™ anti-mouse/human CD45R/B220 (RA3-6B2) | Biolegend | 103227 |
| Pacific Blue™ anti-mouse CD24 Antibody (M1/69) | Biolegend | 101819 |
| Brilliant Violet 711™ anti-mouse CD69 (H1.2F3) | Biolegend | 104537 |
| PE/Cyanine7 anti-mouse CD152 (UC10-4B9) | Biolegend | 106314 |
| APC/Cyanine7 anti-mouse TCR β chain (H57-597) | Biolegend | 109220 |
| Alexa Fluor® 594 anti-mouse CD45.2 (104) | Biolegend | 109850 |
| PerCP/Cyanine5.5 anti-mouse CD45.1 (A20) | Biolegend | 110727 |
| Brilliant Violet 421™ anti-mouse FOXP3 (MF-14) | Biolegend | 126419 |
| PE/Dazzle™ 594 anti-mouse/human Helios (22F6) | Biolegend | 137232 |
| PerCP/Cyanine5.5 anti-mouse CD304 (3E12) | Biolegend | 145208 |
| Alexa Fluor® 488 anti-GFP (FM264G) | Biolegend | 338008 |
| PE/Dazzle™ 594 anti-mouse Ki-67 (16A8) | Biolegend | 652427 |
| Brilliant Violet 421™ anti-mouse CD122 (IL-2Rβ) (5H4) | Biolegend | 105919 |
| PE anti-mouse CD122 (IL-2Rβ) Antibody (TM-β1) | Biolegend | 123210 |
| PE anti-mouse Qa-2 Antibody (695H1-9-9) | Biolegend | 121715 |
| Brilliant Violet 650™ anti-rat CD90/mouse CD90.1 (Thy-1.1) (OX-7) | Biolegend | 202533 |

| | | |
|---|---|---|
| PE/Cyanine7 anti-mouse CD132 (common γ chain) (TUGm2) | Biolegend | 132311 |
| Brilliant Violet 785™ anti-mouse CD45.2 (104) | Biolegend | 109839 |
| Brilliant Violet 421™ anti-mouse CD90.2 (Thy-1.2) (53-2.1) | Biolegend | 140327 |
| PE anti-mouse CD45.1 (A20) | Biolegend | 110708 |
| PerCP anti-mouse CD45 (30-F11) | Biolegend | 103130 |
| PerCP anti-mouse IL-17A (TC11-18H10) | Biolegend | 506944 |
| PerCP/Fire™ 780 anti-mouse CD183 (S18001A) | Biolegend | 155926 |
| APC/Cyanine7 anti-mouse CD62L (MEL-14) | Biolegend | 104428 |
| Purified anti-mouse CD25 Antibody (3C7) | Biolegend | 101902 |
| InVivoMAb anti-mouse CD16/CD32 (2.4G2) | BioXCell | BE0307 |
| InVivoMAb anti-mouse CD8α (2.43) | BioXCell | BE0061 |
| InVivoMAb anti-mouse MHC Class II (I-A/I-E) (M5/114) | BioXCell | BE0108 |
| InVivoMAb anti-mouse CD3ε (145-2C11) | BioXCell | BE0001-1 |
| PE anti-mouse FoxP1 (D35D10) | Cell Signaling Technology | 99657 |
| eBioscience™ APC anti-mouse FOXP3 (FJK-16s) | Invitrogen | 17-5773-82 |
| eBioscience™ PE anti-Phospho-STAT5 (Tyr694) Monoclonal Antibody (SRBCZX) | Invitrogen | 12-9010-42e |
| BioMag Goat Anti-Rat IgG | Qiagen | 310107 |
| Histone H3K27ac (pAb) | Active Motif | 39134 |
| **Chemical, Reagents** | | |
| RBC Lysis Buffer (10X) | Biolegend | 420302 |
| 10% Tween-20 | BioRad | 1662404 |
| Percoll® PLUS | Cytiva | GE17-5445-02 |
| GemCell FBS | GeminiBio | 100-500 |
| 1X PBS pH 7.2 | Gibco | 20012050 |
| HEPES (1 M) | Gibco | 15630106 |
| Sodium Pyruvate (100 mM) | Gibco | 11360070 |
| PBS (10X), pH 7.4 | Gibco | 70011069 |
| GlutaMAX™ Supplement | Gibco | 35050061 |
| RPMI 1640 Medium, no glutamine | Gibco | 21870076 |
| RPMI 1640 Medium, no phenol red | Gibco | 11835030 |
| Digitonin (5%) | Invitrogen | BN2006 |
| Normal Rat Serum | Invitrogen | 01-9601 |
| Proteinase K Solution | Invitrogen | 4333793 |
| Streptavidin | Invitrogen | 434302 |

| | | |
|---|---|---|
| UltraPure™ 0.5M EDTA, pH 8.0 | Invitrogen | 15575020 |
| Brefeldin A | Invitrogen | B7450 |
| BrdU (5-Bromo-2'-Deoxyuridine) | Invitrogen | B23151 |
| Recombinant Human TGF-β1 (CHO derived) | PeproTech | 100-21C |
| Recombinant Mouse IL-2 Protein | R&D Systems | 402-ML |
| DNase I recombinant, RNase-free | Roche | 4716728001 |
| Liberase™ TL Research Grade | Roche | 5401020001 |
| Liberase™ TM Research Grade | Roche | 5401119001 |
| Protector RNase Inhibitor (10000 UNITS) | Roche | 3335402001 |
| Penicillin-Streptomycin | Sigma-Aldrich | P4333 |
| (2-Hydroxypropyl)-β-cyclodextrin solution | Sigma-Aldrich | H5784 |
| Bovine Serum Albumin | Sigma-Aldrich | A2153 |
| Dimethyl Sulfoxide | Sigma-Aldrich | D8418 |
| Dulbecco's Modified Eagle's Medium - high glucose | Sigma-Aldrich | D5796 |
| Ionomycin calcium salt | Sigma-Aldrich | I0634 |
| Phorbol 12-myristate 13-acetate | Sigma-Aldrich | P8139 |
| Tamoxifen | Sigma-Aldrich | T5648 |
| Paraformaldehyde, 16% w/v | Alfa Aesar | 43368 |
| 2-Mercaptoethanal | Sigma-Aldrich | 63689 |
| **Commercial assays** | | |
| D1000 Ladder | Agilent | 5067-5586 |
| D1000 Reagents | Agilent | 5067-5583 |
| D1000 ScreenTape | Agilent | 5067-5582 |
| TaqMan™ Universal Master Mix II, with UNG | Applied Biosystems | 4440042 |
| BD Pharmingen™ APC BrDU Kit | BD biosciences | 552598 |
| SPRIselect, 5 mL | Beckman Coulter | B23317 |
| Zombie Aqua™ Fixable Viability Kit | Biolegend | 423102 |
| Zombie NIR™ Fixable Viability Kit | Biolegend | 423105 |
| Dynabeads™ Mouse T-Activator CD3/CD28 for T-Cell Expansion and Activation | Gibco | 11456D |
| CellTrace™ Violet Cell Proliferation Kit, for flow cytometry | Invitrogen | C34557 |
| CountBright™ Absolute Counting Beads | Invitrogen | C36950 |
| eBioscience™ Annexin V Apoptosis Detection Kits PE | Invitrogen | 88-8102-72 |
| eBioscience™ Foxp3 / Transcription Factor Fixation/Permeabilization Concentrate and Diluent | Invitrogen | 00-5521-00 |
| eBioscience™ Permeabilization Buffer (10X) | Invitrogen | 00-8333-56 |

| | | |
|---|---|---|
| GFP BrightComp eBeads™ Compensation Bead Kit | Invitrogen | A10514 |
| CellEvent™ Caspase-3/7 Detection Reagents, Red, Powder | Invitrogen | C10430 |
| UltraComp eBeads™ Compensation Beads | Invitrogen | 01-2222-42 |
| SuperScript™ III First-Strand Synthesis System | Invitrogen | 18080051 |
| ArC™ Amine Reactive Compensation Bead Kit | Invitrogen | A10346 |
| NEBNext® High-Fidelity 2X PCR Master Mix | NewEngland Biolab | M0541S |
| MinElute PCR Purification Kit (50) | Qiagen | 28004 |
| RNeasy Micro Kit (50) | Qiagen | 74004 |
| **TaqMan probes** | | |
| *Actb* | Thermo Fisher | Mm02619580_g1 |
| *Foxp1* | Thermo Fisher | Mm01181991_g1 |
| *Foxp2* | Thermo Fisher | Mm00475030_m1 |
| *Foxp3* | Thermo Fisher | Mm00475162_m1 |
| *Foxp4* | Thermo Fisher | Mm01269228_g1 |
| *Il2ra* | Thermo Fisher | Mm01340213_m1 |
| **Animals** | | |
| $Foxp3^{tm9(EGFP/cre/ERT2)Ayr}/J$ | Jackson Laboratory | 016961 |
| $Foxp1^{fl/fl}$ | from H. Tucker | Feng et al.(*9*) |
| $Foxp4^{fl/fl}$ | from E. Morrissey | Li et al.(*42*) |
| $Foxp3^{YFP/cre}$ | from A. Rudensky | Rubstov et al.(*43*) |
| $Cd4^{Cre/ERT2}$ | from F. Gounari | Aghajani et al.(*52*) |
| $B6.129X1\text{-}Gt(ROSA)26Sor^{tm1(EYFP)Cos}/J$ | from F. Costantini | Srinivas et al. (*44*) |
| $B6.PL\text{-}Thy1^{a}/CyJ$ | Jackson Laboratory | 000406 |
| *B6.SJL-Ptprca Pepcb/BoyJ* | Jackson Laboratory | 002014 |

| | | |
|---|---|---|
| *B6.129P2-Tcrb$^{tm1Mom}$ Tcrd$^{tm1Mom}$/J* | Jackson Laboratory | 002122 |
| **Software and algorithm** | | |
| Flow Jo™ (Ver10.10) | BD Biosciences | https://www.flowjo.com |
| GraphPad Prism (Ver10.2.2) | GraphPad Software | https://www.graphpad.com/ |
| R (Ver4.5.0) | R Foundation | https://www.r-project.org |
| RStudio | Posit Software | https://posit.co |
| Docker | Docker Inc. | https://www.docker.com |
| IGV | UC San Diego | https://igv.org |
